# Programming Microbe-Computer Interaction for Customized Life-Nonlife and Cross-Life Communication

**DOI:** 10.1101/2025.09.27.679003

**Authors:** Zhijie Feng, Chao Zhang, Xin Huang, Yingying Zhang, Hui Liu, Meihui Cui, Hao Zhang, Zihe Song, Mengya Li, Hongxiang Li, Binglin Ma, Shenjunjie Lu, Mingshan Li, Guijie Bai, Dawei Sun, Xinyu Zhang, Zhenping Zou, Ying Zhou, Bangce Ye, Miao Gao, Huifang Yin, Minzhen Tao, Peiyuan Liu, Duo Liu, Haibin Luo, Hanjie Wang

**Affiliations:** Key Laboratory of Tropical Biological Resources of Ministry of Education and Hainan Engineering Research Center for Drug Screening and Evaluation, School of Pharmaceutical Sciences, Hainan University, Haikou 570228, China; School of Life Sciences, Faculty of Medicine, Tianjin University, Tianjin 300072, China; State Key Laboratory of Synthetic Biology, Tianjin University, Tianjin 300072, China; School of Medical Imaging, Xuzhou Medical University, Xuzhou, 221004, China; Thoracic Surgery Laboratory, Xuzhou Medical University, Xuzhou, 221006, China; Laboratory of Biosystems and Microanalysis, State Key Laboratory of Bioreactor Engineering, East China University of Science and Technology, Shanghai 200237, China; School of Marine Science and Technology, Tianjin University, Tianjin 300072, China; Engineering Research Center for the Prevention and Control of Animal Original Zoonosis of Fujian Province University, College of Life Science, Longyan University, Fujian 364012, China

## Abstract

This extension of human-computer interaction (HCI) paradigm to the non-intelligent microbe-computer interaction (MCI) is attractive for building generalized life-nonlife communication. Main challenges lie in how to construct biohybrid interface within individual MCI unit and thereafter program cross-life communication based on these MCI units. Here, we propose the design and construction of opto-interfaced MCI unit composed of engineered microbes and computer terminals, for enabling quantifiable transduction between microbial signals and digital signals. These units were able to serve as biohybrid sensors, actuators and transceivers, supporting biomonitoring, bio-actuation and microbe-centric bidirectional operations. Based on these functional units, we established programmable cross-life communication architectures capable of unicast, multicast, and logic-gate-driven communication with temporal dynamic control. The architectures were validated in biomedical scenarios including tumor hypoxia biomarking, epidermal heavy metal detoxification, and colitis intervention, demonstrating robust performance in closed-loop diagnostic-therapeutic applications. Additionally, we integrated MCI with an unmanned surface vessel for environmental monitoring, highlighting its cross-domain potential. Collectively, we provide a scalable strategy for building customized life-nonlife and cross-life communication, which is expected as an important complement to internet of things (IoT).

## Introduction

The development of human-computer interaction (HCI) has yielded substantial benefits for human society, such as lowering barriers to machine usage, improving health and social experiences, and deepening the connection to internet of things (IoT)^1^. With the deep integration of biotechnology (BT) and information technology (IT), the interactive objects of computers are extending from cognitively capable humans to generalized non-intelligent biosystems, which is expected to achieve living biosystem-computer interaction (LBCI)^2–5^. The LBCI aims not merely to create individual biohybrid units, but to complement the IoT where the interaction of living biosystems, artificial intelligence (AI) and mechanic devices will benefit health^6–8^, environment^9^, agriculture^10^, biohybrid robotics^11^ and intelligence^12^, etc. A key point of LBCI is to establish life-nonlife information flows at the biohybrid interface, where the human-centric input commands in HCI shall be shifted into the sensing and actuation of particular life-centric functions in LBCI^3,4,13^. The high diversity and heterogeneity of multicellular organisms make decoupling and quantifying information flows exceedingly difficult. Given this challenge, unicellular microbes with simplified and more predictive biological processes and communities are priority^14,15^. The establishment of microbe-computer interaction (MCI) will contribute to the future more complex LBCI.

The primary task to establish MCI is to realize life-nonlife signal transduction and information flow at the biohybrid interface. On the first hand, diverse studies have programmed microbes with tools of genetic circuits^16–19^, genome synthesis^20,21^ and editing^22,23^ to realize transduction between analog signals of biological functions and quantifiable biochemical or biophysical signals. On the other hand, the integrated circuits of electronic elements as computer terminals can record or trigger real-world signals with digital signals in a programmable way. Thus, the analog-digital signal transduction at MCI interface is achievable. Pioneering studies include Lu group’s microbial-electronic capsules to transduce bioluminescent signals into photocurrent signals^6,7^ and Ajo-Franklin group’s microbial-electrodes to transduce bioelectrical signals into measurable electrical signals^24^. Besides, digital forms of light^25^, electricity^26^, sound^27^ and magnetic^28^ signals have been utilized to activate preset biological functions. Our group has established bidirectional signal transduction between engineered microbes and electronic capsules with the opto-interfaced signals^8^. In general scenarios where different microbial populations and even other organisms are involved, it is necessary to connect individual MCI units to construct cross-life communication^29–31^. This will expand the boundaries of human cognition and control, benefiting applications in many fields.

The necessity of MCI-based cross-life communication is to expand the constrained distance, accuracy and latency of communication across cell communities relying on biochemical signals^32,33^. MCI-based communication is expected to overcome the spatiotemporal constraints to build connection among distant cell communities. This process is possible in theory but there are still practical problems to be solved. A first problem is how to connect MCI units to achieve controllable architecture of cross-life communication. Especially, the MCI-based communication shall realize remote point-to-point accuracy, logic programmability, temporal tunability, and meanwhile reserve human directive channel to achieve recording, training and debugging. Secondly, the applicable benefits of such MCI-based communication are ready to explore and validate. For instance, in biomedical scenarios, it is worthy to establish a wearable, portable, ingestible or some else form of MCI-based communication to support convenient disease biosensing and bio-intervention presently relying on specialized medical instruments and stuffs. In environmental scenarios, it is worthy to build an unmanned and AI-supported form of MCI-based communication to support active environmental hazard monitoring and remediation. MCI-based communication is expected to facilitate more extensive scenarios in future.

This study proposes a strategy of programming MCI for life-centric and cross-life communication. The optical signals of tunable intensity and spatiotemporal resolution are utilized to build contact-independent MCI interface, generating customized MCI unit. These MCI units are capable of transducing biological signals into computer-coped electronic signals or vice versa in quantifiable modes, serving as microbe-to-computer unit (MCU) or computer-to-microbe unit (CMU) (Figure 1a). Further, we connect these units in assistance of Bluetooth to program various cross-life communication architectures, such as unicast, multicast and logic-gate cast, achieving controllable directionality, logics and dynamics of information flows (Figure 1a). Ultimately, we demonstrate several application examples, including tumor identification and intervention, epidermal heavy metal ion sensing and chelation, intestinal inflammation diagnosis and alleviation, and even unmanned surface vessel (USV)-based aqueous sampling and analysis. Our proposed strategy holds transformative potential in multiple fields, such as the sophisticated design and manipulation of biohybrid systems, personalized and precision healthcare, intelligent and autonomous environmental monitoring, and other aspects related to cross-life connection and collaboration.

**Fig. 1.**
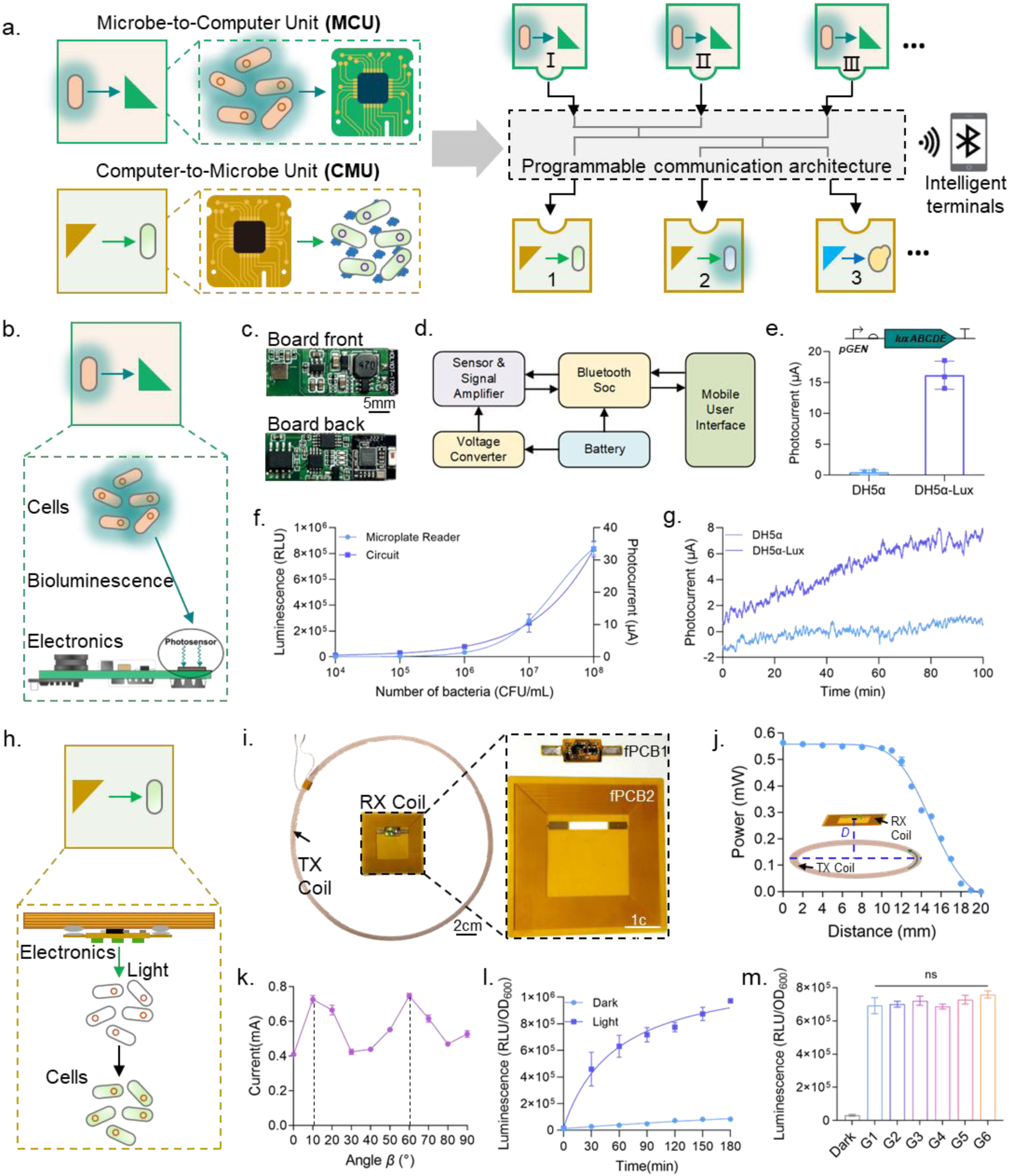
The concept of MCI-based communication and construction of MCI unit models. **a)** Establishment of programmable architecture MCI-based communication with MCU and CMU. **b)** Schematic diagram of bioluminescence-to-photocurrent signal transduction in MCU. **c)** Image of the integrated electronic circuit board in MCU. **d)** Circuit design schematic of the integrated board. **e)** Photocurrent measurements of luminescent strain (DH5α-Lux) by MCU. **f)** Quantitative detection of luminescence density and CMU-based photocurrent in DH5α-Lux cultures. **g)** Continuous monitoring of luciferase expression in DH5α-Lux via CMU. **h)** Diagram of LED-based optical activation for optogenetically engineered microbial functions using CMU. **i)** Image of the rechargeable CMU unit composed of three parts: TX Coil (transmitter coil), fPCB1 (LED circuit), and fPCB2 (receiver coil). **j)** Measured power output of the electronic module at different distances between receiver and transmitter coils. **k)** Effect of the angle between enhancement coil and transmitter coil on current in the receiver coil. **l)** Time-dependent activation of luciferase expression in optogenetically engineered EcN under green light irradiation via CMU. **m)** Luciferase activation in microbial cultures at different distances between receiver and transmitter coils; G1 to G6 correspond to 10, 12, 13, 14, 15, and 19 cm spacing. * 0.01<p<0.05, ** 0.001<p<0.01, *** p<0.001.

## Results

### Construction of Opto-Interfaced MCI Units

In order to realize microbe-computer interaction (MCI), a pivotal task is to establish quantifiably conversion between bio-specific signals and electronic signals. Thus, we designed and constructed two basic units for opposite direction of signal transduction, namely microbe-to-computer Unit (MCU) and Computer-to-Microbe Unit (CMU). Both units were composed of engineered microbes and computer terminals (e.g. integrated semiconductors, smartphones, etc.). The interface within MCU was designed as a contact-independent pattern, where the bacterial bioluminescent signal was chosen to overcome strict contact-dependence for bioelectronic signal transduction (Fig. 1b). To showcase the MCU function, we constructed a primary model containing bioluminescent microbial cells and integrated electronic circuit board with merely a cover glass flat interface, for the transduction of bioluminescent signals into electrical signals (Fig. 1c). The circuit board consisted of an optimized photosensor for bioluminescence-to-electrical signal conversion, a signal amplifier, and auxiliary functional components (Fig. 1d, S1a). A smartphone application (APP) was developed for monitoring the real-time photocurrent values detected by the unit (Fig. S1b-e). After coped by a data filtering algorithm, the photocurrent values were recorded without affecting the average value within any 1-minute window, but the dynamic tendency was precisely preserved (Fig. S1f, S1g).

We tested a series of conditions to verify the functionality and quantitative rules of the MCU model. Under darkness, detection of the control strain and luciferase-expressing strain (DH5α-Lux) by MCU got significantly divergent photocurrent values (Fig. 1e). As bioluminescent bacterial density increased exponentially, a strong positive correlation persisted between microplate reader-measured fluorescence intensity and our MCU-recorded photocurrent (Fig. 1f, Fig. S1h). During proliferation in culture medium, our MCU accurately reflected dynamic changes in total luciferase expression while remaining unresponsive to non-luminescent cell density (Fig. 1g). Beyond *E. coli*, the MCU model was also able to detect the bioluminescence of other organisms including bacteria (*Bacillus subtilis*), fungi (*Saccharomyces cerevisiae*), and mammalian cells (HEK293T cells), demonstrating broad applicability (Fig. S2a-d). To simulate a real-world scenario in which biological systems might approach a computer before interacting with it, we set up two models. In a first model, we encapsulated the electronic board within a light-transmissive capsule and successfully detected the bioluminescent bacteria (*E. coli* Nissle 1917, EcN) adhering to porcine intestinal segments *in vitro* during capsule transit (Fig. S2e-g). In a second model, we configured the electronic board in a light-proof box containing microneedle array enabling negative pressure sampling^34^ (Fig. S3). The bioluminescent bacteria cells externally approaching to form complete MCU were successfully detected. These models were customized forms of MCU.

For reversal transduction of electrical signals to biological signals, the CMU was constructed by combining computer-controlled light sources (e.g. laser, light-emitting diode (LED), etc.) and optogenetically engineered biosystems. To showcase CMU function, we constructed a model containing optogenetically engineered EcN-Opto-LuxA and micro-LED controlled by a rechargeable integrated circuit (Fig. 1h). This CMU leveraged contactless power delivery from the transmitting (TX) coils to the receiving (RX) coils in the form of two-layer flexible printed circuit board (fPCB) integrated with micro-LEDs (Fig. 1i). The effects of coil distance and angular alignment on charging efficiency were systematically examined. The results demonstrated that the full power transfer was achieved within a 12 mm separation distance and at the angular offsets of either 10° or 60° relative to the TX coil (Fig. 1j, 1k). Subsequently, we evaluated the effects of observed power attenuation over distance on the activation of optogenetically engineered bacterial functions. The activated GFP expression exhibited a time-dependent increase under continuous LED irradiation, approaching saturation over extended periods (Fig. 1l). Furthermore, the green-light-responsive EcN strain were subjected to the LED irradiation in fPCB powered by TX coils at different power transfer distances (12-19 cm), showing that even the marginal power transfer at 19 cm was sufficient to induce substantial GFP reporter expression (Fig. 1m). These results showed that the CMU enabled transduction of electronic signals into bio-activating signals, establishing a functional information flow. Collectively, we established several plug-and-play MCI toolkits for supervising microbe-centric functions.

### Expansion of MCI Units as Biohybrid Sensors, Actuators and Transceivers

The customized forms of MCU and CMU shall not only monitor or trigger biosystem internal signals, but also interact with external living world, serving as biohybrid sensors, actuators and transceiver. Thus, firstly, we tested the MCU performance of sensing and transducing external biomolecule signals (Fig. 2a). We fabricated a light-proof box via 3D printing as a MCU sensor configuration model, where the bacteria engineered with biosensing genetic circuits were housed in (Fig. 2b, Fig. S4a, Fig. S4b). External biomolecule sample introduction enabled photocurrent measurements. Initial tests with varying L-arabinose concentrations demonstrated high correlation between photocurrent and luminescence detection values (Fig. 2c). The MCU sensor further generated dynamic time-dependent response profiles to certain fixed arabinose concentration (Fig. 2d). To assess versatility, we incorporated the engineered EcN responsive to distinct biomolecules (1 mM nitrate^8^, 100 μg/L tetracycline^35^ and 1 μM heme^36^) into the respective MCU (Fig. 2e, S4c-e). Significant photocurrent differentials emerged between induced and non-induced bacterial wells after respectively sufficient incubation periods. Additionally, by spiking tetracycline of graded concentrations into murine whole blood samples, we achieved dose-dependent photocurrent responses using our MCU, confirming functionality in complex biological matrices (Fig. S4f, S4g). At present stage, we expanded the MCU function as modular sensors combining both the biological responsive specificity of microbes and the low latency of electronics.

**Fig. 2.**
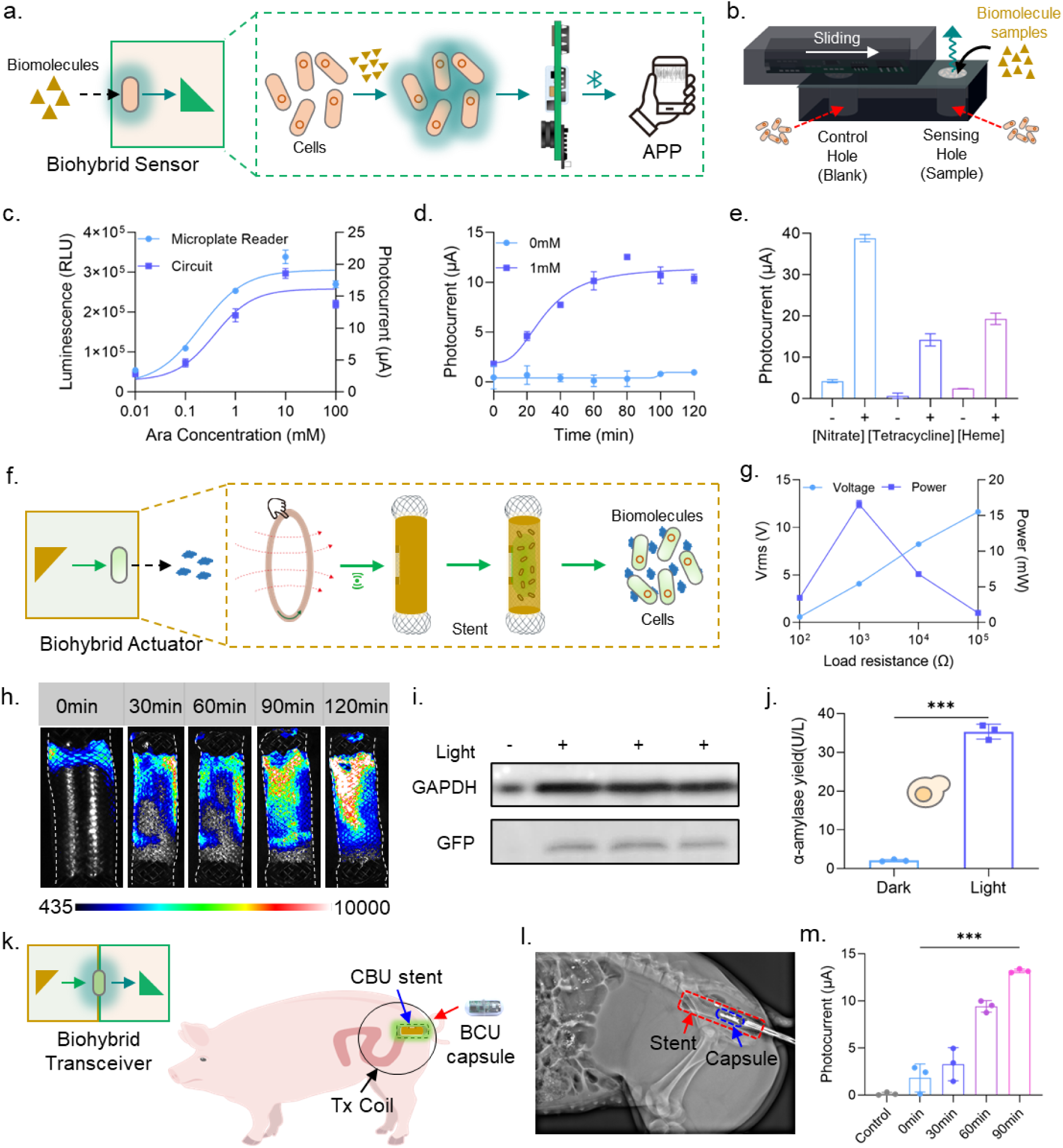
Evaluation of expanded MCI units as biohybrid sensors, actuators, transceivers. **a)** Schematic of the MCU-based biohybrid sensor. **b)** Physical embodiment of the MCU sensor in a light-proof box. **c)** Photocurrent detected by the MCU sensor and luminescence density detected by a microplate reader for the EcN sensing different L-Arabinose concentrations. **d)** Time-dependent monitoring of L-Arabinose by the MCU sensor. **e)** Sensing of various biomolecules using the MCU. **f)** Diagram of a CMU-based biohybrid actuator mounted on a 3D stent. **g)** Electrical response of the stent-mounted CMU under different loading conditions. **h)** Spatiotemporal dynamics of luciferase expression in EcN induced by green light via the CMU stent. **i)** Western blot analysis of GFP secretion from EcN under green light induction controlled by the CMU stent. **j)** Assay of amylase secretion from *S. cerevisiae* induced by blue light via the CMU stent. **k)** Diagram of a biohybrid transceiver formed by a CBU stent and BCU capsule in porcine intestine for simultaneous actuation and feedback monitoring of EcN. **l)** X-ray image of the transceiver in the porcine rectum. **m)** Control and monitoring of engineered EcN function in the porcine intestine using the transceiver. * 0.01<p<0.05, ** 0.001<p<0.01, *** p<0.001.

Further, we tested the CMU performance as a spatiotemporally controllable actuator. We engineered a three-dimensional stent^37^ with a special barrel shape to house both the RX coil and optogenetically engineered bacteria (Fig. 2f). This deformable stent permitted up to 50% compressive or tensile deformation while allowing shape recovery (Fig. S5a). Subsequently, the LED components and RX coil-embedded fPCB were assembled onto the stent, yielding a stable integrated construct (Fig. S5b). Testing confirmed that the higher carrying capacity could be achieved within appropriate load range (Fig. 2g). The stent’s presence exhibited no detectable impact on the RX coil’s open-circuit voltage (Fig. S5c), while the current output remained stable at bending angles below 280° (Fig. S5d). In addition, *in vitro* porcine tissue was employed to test charging features (Fig. S5e). Evaluations of compression ratios and angular alignments relative to the TX coil demonstrated the reliable operation across all tested conditions and the optimal operational parameters (Fig. S5f, S5g). To assemble the integrated stent-form CMU actuator, biocompatible polyvinyl alcohol (PVA)^38^ was selected for packaging the green-light-responsive EcN onto the stent surface, due to its negligible impact on bacterial viability and growth kinetics (Fig. S6a-c). Over 120-minute irradiation in the CMU actuator, there exhibited obviously enhanced luminescence intensity, suggesting bacterial luciferase expression (Fig. 2h). The secretion of another reporter, green fluorescent protein (GFP), was also verified under the control of CMU actuator (Fig. 2i). We also validated the CMU’s capability in controlling α-amylase synthesis and secretion in optogenetically engineered yeast (Fig. 2j, S6d, S6e). Collectively, we established a wirelessly rechargeable CMU actuator model to control microbes, enriching the integration-split deployment forms and the enhancing accuracy of spatiotemporal control.

When combing MCU sensor and CMU actuator, we boldly imagined whether a life-centric transceiver could be realized to make each microbial system as an information transmission node. This was easily imagined to achieve *in vitro*. Thus, we decided to assemble the transceiver in a highly complex *in vivo* environment. Through transanal catheter deployment, the stent-form CMU actuator was successfully implanted within the porcine rectum, demonstrating stable positional retention at targeted sites (Fig. S7a). Externally controlled LED activation was reliably achieved (Fig. S7b-c), with histopathological analysis confirming an absence of inflammation or tissue damage (Fig. S7e, S7f). To validate real-time functional protein synthesis in the green-light-responsive EcN during sustained illumination, the capsule-form MCU built above was loaded to the CMU location by a similar transanal way (Fig. 2k, 2l, S7g). The combination of CMU actuator and MCU sensor was proved to enable continuous monitoring of the dynamically controlled expression of bacterial luciferase *in vivo*, generating a microbe-centric transceiver (Fig. 2m). So far, we built and verified varying functional biohybrid sensors, actuators and transceivers, remarkably enhancing the modularity, reusability, deployability and accuracy of MCI.

### Establishment of MCI-Based Cross-life Communication Architecture

Based on the above biohybrid sensors, actuators and transceivers, we explored whether the cross-life communication could be realized. The key point was to test the remote point-to-point effectiveness, logic architecture programmability and temporal tunability of cross-life communication. Initially, we verified a fundamental unicast communication model to generate information flow from a sender strain to a receiver strain (Fig. 3a). MCU sensor was used to transduce the bioluminescence signal of DH5α-Lux (sender strain) to photocurrent signal under dark conditions (Fig. 3b). When this photocurrent value exceeded the preset 6 μA threshold, the MCU emitted a wireless signal that successfully activated another CMU actuator containing green-light-responsive strain EcN-Opto-GFP (receiver strain) through the Bluetooth signal transmission controlled by smartphone APP. Thereby, the reporter GFP expression in the receiver strain was significantly triggered and increased after 40 min (Fig. 3b). In addition, we validated a multicast communication model enabling sender-to-double-receiver information transmission (Fig. 3c), where the processing of single bioluminescence input signal successfully triggered two distinct output signals in the form of target protein expression (Fig. 3d). Collectively, these experiments confirmed the remote point-to-point effectiveness of MCI-based communication.

**Fig. 3.**
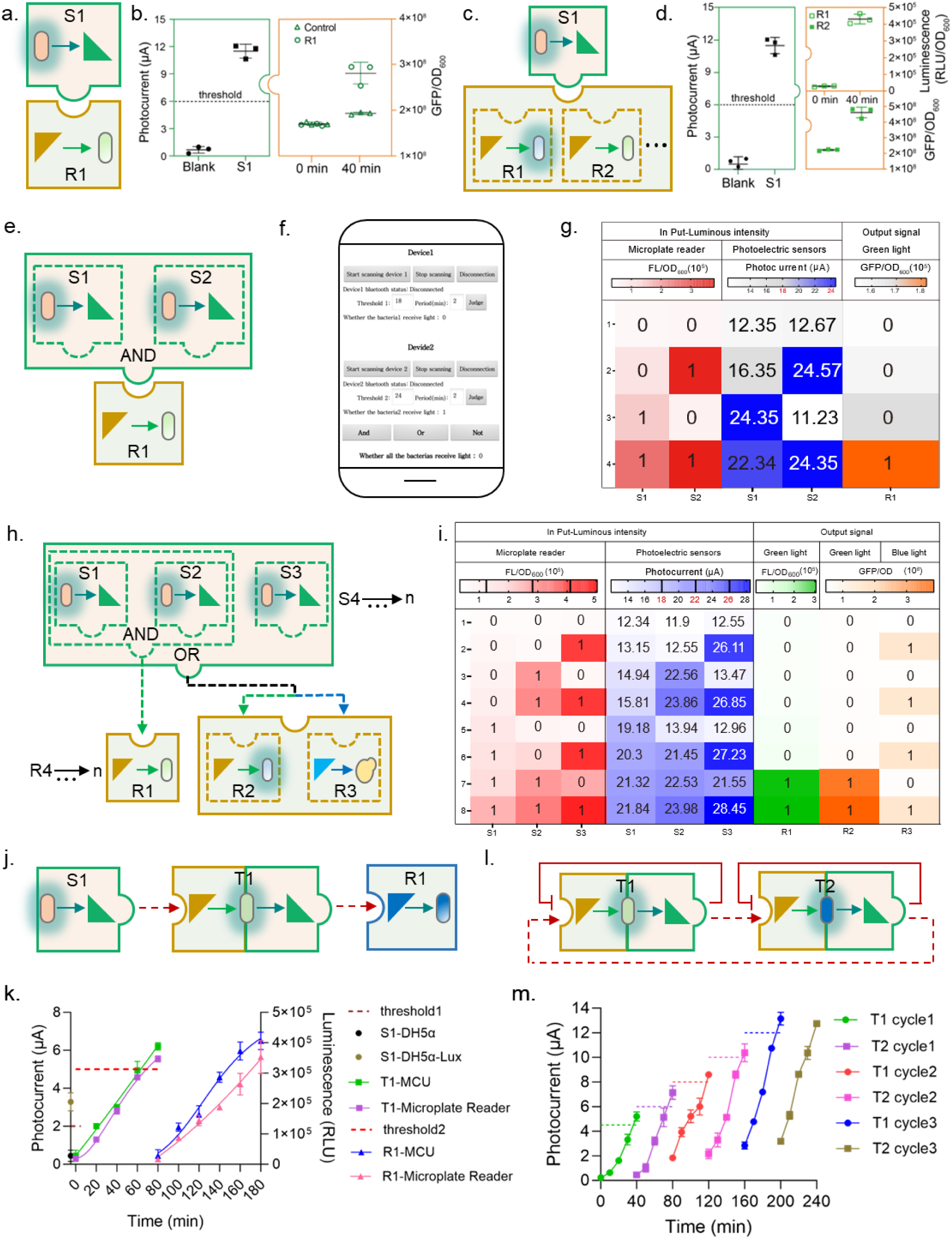
Establishment of MCI-based cross-life communication architectures. **a, b)** Establishment and test of unicast communication between a sender microbial community (S1, MCU) and a single receiver microbial community (R1, CMU). **c, d)** Establishment and test of multicast communication from a single sender (S1) to two distinct receivers (R1 and R2). **e)** Establishment of AND-gate logic communication involving two senders (S1, S2) and one receiver (R1). **f)** A smartphone application (APP) enabled user-friendly programming of the activation thresholds for S1 and S2, and also supports diverse logic operations (e.g., OR, NOT gates). **g)** Characterization of the AND-gate communication output (R1 activity) for all four possible input combinations of S1 and S2. h) Design of a cross-life communication mode implementing three senders and three receivers with more multi-level logic gates. **i)** Characterization of the output results of receivers for all eight possible input combinations from three senders. **j, k)** Diagram and test of a cascaded communication chain across three sequentially connected microbial communities. k) Characterization of the cascaded chain. **l)** Design of feedback communication loop established between two microbial communities as transceivers (T1, T2), which could be programmed to implement either positive or negative feedback. **m)** Characterization of the positive (or negative) feedback communication loop.

Building upon this foundation, we further investigated the logic architecture programmability of MCI-based communication. We established an AND-gate communication mode to determine the output signal according to the logic calculation results of the two inputs (Fig. 3e). This logic calculation was programmed in APP (Fig. 3f). For instance, we set the photocurrent threshold for sender strain 1 at 18 μA and sender strain 2 at 24 μA. Only when both thresholds were reached would the AND-gate program on the APP activate, subsequently triggering the optogenetic function in the receiver strain. Experiments confirmed the four different AND-gate results (Fig. 3g). Additionally, the APP interface allowed convenient program of OR-gate, NOT-gate, and other logic operations. To demonstrate the diversity of logic architecture, we implemented a multiple-senders-to-multiple-receivers communication mode (Fig. 3h). Photocurrent thresholds for senders 1, 2, and 3 were set at 18 μA, 22 μA, and 26 μA, respectively. When senders I and II simultaneously emitted positive signals, an AND-gate operation activated receiver 1. This AND-gate output meanwhile also underwent an OR-gate operation with sender III’s signal, ultimately controlling the biological functions of EcN receiver 2 and yeast receiver 3. Experimental validation confirmed the precise execution of all the eight possible input-output results according to predefined logics (Fig. 3i, S8a-c). To sum up, these experiments confirmed the programmability of MCI-based communication.

Ultimately, the temporal tunability of MCI-based communication was evaluated. We tested the two communication models, namely cascaded and dynamically regulated feedback modes. For the cascaded mode, a sender activated a transceiver, which subsequently triggered the final biological output of a receiver (Fig. 3j). Experimental results demonstrated successful cascaded signal propagation (Fig. 3k). Only when the supervised photocurrent of transceiver reached the preset threshold of 5 μA, would the receiver function be triggered. Regarding dynamically regulated feedback mode, we programmed two transceivers 1 and 2 with cross-life positive feedback along with each self-feedback inhibition for the purpose of building cross-life oscillatory pattern (Fig. 3l). It was shown that once one of the transceivers reached the preset threshold, it successfully triggered self-shutting-off and activated the function of the other transceiver (Fig. 3m, S8d). Above all, these experiments confirmed the temporal tunability of MCI-based communication. Besides the above, more diverse communication models were probable to be realized, including feedback or feedforward loops, the hybrid models also containing biochemical-based short-range communication (Fig. 3m, S8e, S8f) and the networked models integrated with Local Area Network (LAN) and internet (Fig. S8f), which would greatly enrich life-centered and cross-life information exchange capabilities. Collectively, we systematically propose the concept, basic construction mode and scalability of MCI-based cross-life communication.

### Program MCI-based communication for tumor biomarking and intervention

After validating several functional models of MCI-based communication, we sought to test its applicability in biomedical scenarios^39^. We configured the sensing function of the MCU sensor to detect hypoxic conditions^40^, mimicking the local tumor microenvironment, thereby driving the CMU actuator to synthesize and produce a PD-L1 nanobody (PD-L1nb) capable of exerting tumor immunotherapy effects (Fig. 4a). Results showed that under *in vitro* aerobic (conventional ambient conditions, 21% O₂) and hypoxic conditions (below 0.5% O₂) similar to tumor microenvironment, the MCU sensor containing hypoxia-sensing bacteria exhibited a significant difference in both luminescence and photocurrent values (Fig. 4b). Furthermore, triggered wirelessly by this result, the green light-irradiated bacterial strain within the CMU actuator synthesized and secreted a markedly different quantity of PD-L1nb (Fig. 4c). This experiment demonstrated that the MCU and CMU successfully cooperated *in vitro* to establish efficient unicast communication, realizing the complete information flow from hypoxia detection to nanobody production.

**Fig. 4.**
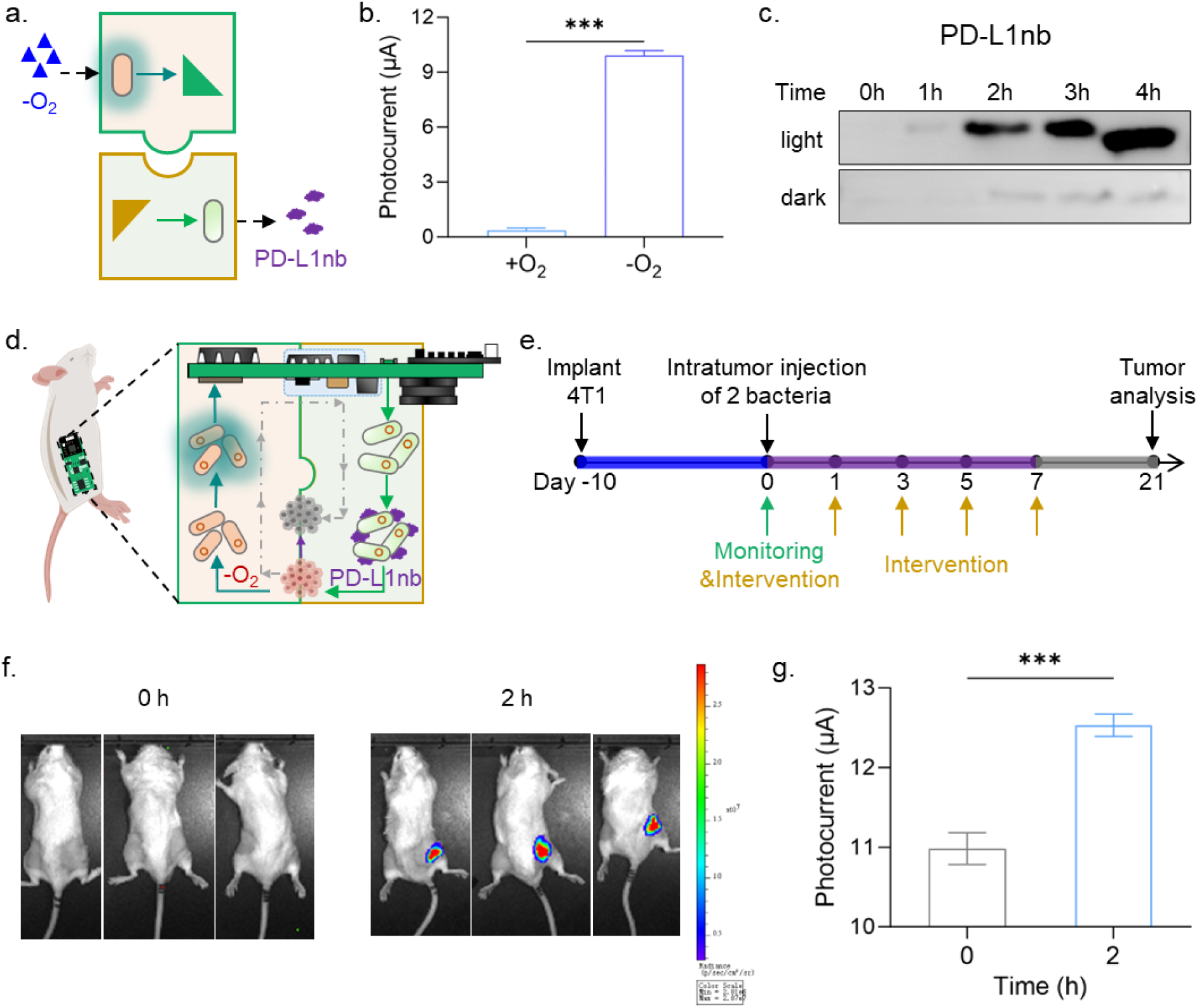
Evaluation of a wearable form for MCI-based unicast communication for tumor biomarking and intervention. **a)** Diagram of MCI-based unicast communication where an MCU sensing the tumor-microenvironment hypoxic would trigger another CMU to express and release nanobody PD-L1nb. **b)** Characterization of the MCU for sensing hypoxic conditions. **c)** Characterization of the CMU for light-controlled expression and secretion of PD-L1nb. **d)** Diagram of the unicast communication system implemented via a wearable device and local injection of engineered bacteria in mice for tumor biomarking, intervention, and whole-process supervision. **e)** Timeline of the monitoring of developed tumor and therapeutic intervention in mice. **f)** Characterization of hypoxia-sensing bacterial function in the tumor region using an imaging system. **g)** Photocurrent-based measurement of hypoxia-sensing bacterial function in the tumor region via the MCU. * 0.01<p<0.05, ** 0.001<p<0.01, *** p<0.001.

Subsequently, we established an *in vivo* mouse model for tumor recognition and intervention to test the functionality of a wearable device based on the principle of MCI-based unicast communication (Fig. 4d). A timeline was designed wherein 4T1 tumor cells were implanted into the flank of mice 10 days before the start of the experiment. Starting from day 0, a mixture of sensing bacteria (DH5α-Hypoxia) and therapeutic bacteria (EcN-Opto-secPD-L1nb) was injected into the tumor, followed by additional injections of therapeutic bacteria on days 1, 3, 5, and 7 (Fig. 4e). The integrated circuit board worn by the mice integrated both MCU sensor and CMU actuator functionalities, utilizing a shared wireless communication module to transmit and receive information via Bluetooth to a smartphone for processing within an APP interface. Experiment results showed that, compared to aerobic conditions, the DH5α-Hypoxia injected simultaneously into tumor-bearing mice exhibited a significant enhancement in both bioluminescence and photocurrent (Fig. 4f, 4g). The bioluminescence intensity at the tumor site increased initially, peaking at 2 hours post-injection, before gradually decreasing. The reduction in bioluminescence resulted from the limited availability of the fatty aldehyde, a chromophore synthesized in oxygen-dependent pathway for the bacterial luciferase (Lux).

Further, the MCU photocurrent over preset threshold triggered the CMU to initiate green light irradiation for 4 hours every other day. This programmed instruction activated the microbial PD-L1nb expression and secretion to block the PD-L1 immune checkpoint. To sum up, we established MCI-based communication mode across the engineered biosensor microbes and biotherapeutic microbes in the hypoxic core of tumors. This mode could provide tumor biomarking and intervention functions, serving as an important supplement to existing tumor labeling and therapeutic strategies^41–43^. In contrast to previously reported self-tunable biotherapeutic strategies, our mode remained under human supervision and control, while supporting future networked data management and decision-making via smart terminals. As a miniaturized and portable device, this wearable MCI-based communication modality was expected to complement imaging methods that rely on expensive instrumentation, offering a reusable and cost-effective approach to long-term health monitoring.

### Program MCI-based communication for epidermal and intestinal healthcare applications

To expand the application potential of MCI-based communication, we investigated its utility in epidermal and intestinal healthcare. First, we designed a model in epidermal scenario for MCU-based biosensing and CMU-based bio-chelation of the potentially contacted heavy metal ions (e.g. Pb²⁺) (Fig. 5a). Using a mouse model, this system simulated a scenario in which a population was exposed to the same hazardous heavy metal ion (e.g. Pb^2+^) environment. The positive detection signals on individual skin would trigger the activation of MtnB protein secretion across the population for Pb^2+^ chelation (Fig. 5b). We conducted a series of *in vitro* and *in vivo* experiments. First, EcN bacteria equipped with a Pb²⁺-sensing genetic circuit^44^ (Fig. S9a) were used to detect Pb²⁺ solutions. Results showed a strong positive correlation between both luminescence intensity and MCU-measured photocurrent values with increasing Pb²⁺ concentrations (Fig. S9b, S9c). We further evaluated whether CMU-activated expression of MtnB in EcN could chelate Pb²⁺ (Fig. S9d). Decreases in both luminescence and photocurrent results indicated successful reduction of free Pb²⁺ (Fig. S9e, S9f). We then simulated epidermal Pb²⁺ sampling and detection by applying and air-drying droplets of Pb²⁺ solutions (0, 10, 20, 50 μM) on a petri dish surface (Fig. S9g). Both luminescence and MCU-measured photocurrent values exhibited a positive correlation with Pb²⁺ concentration (Fig. S9h, S9i). Finally, we tested the mouse model. A preliminary application of 10 μM Pb²⁺ solution on the mouse skin allowed us to establish an MCU-measured threshold photocurrent value of 6 μA (Fig. S9j, 5c). The population of mice were fitted with annular patches containing hydrogel dressings loaded with the light-controlled EcN bacteria designed to secrete MtnB (Fig. S9k). An integrated circuit board with LEDs was attached around the patch (Fig. 5d), forming a complete CMU device carried by each mouse (Fig. 5e). When an individual mouse yielded a positive signal exceeding the 6 μA threshold, a smartphone APP remotely activated the CMUs across the population via Bluetooth. Although direct validation of Pb^2+^ chelation on actual skin remained challenging, we confirmed the expression, secretion and epidermal retention of reporter protein (mCherry) upon CMU activation after removing the wearable CMU (Fig. S9l, 5f). These results demonstrate the feasibility of MCI-based communication for epidermal healthcare maintenance using biological means, serving as an necessary supplement to the epidermal care field.

**Fig. 5.**
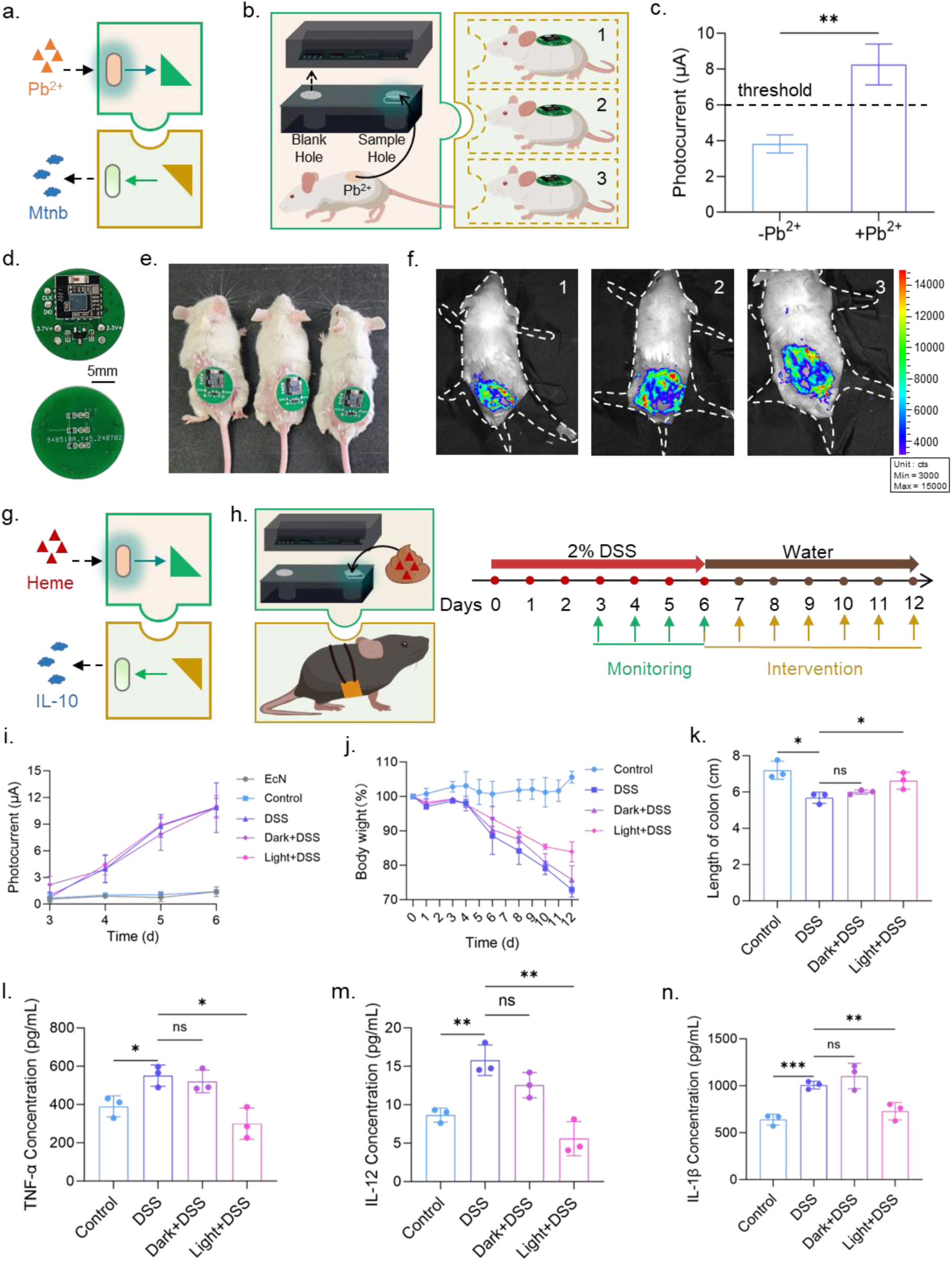
Communication between box-form MCU and wearable CMU for monitoring-driven intervention in epidermal and intestinal scenarios. **a)** Diagram of the MCU sensing heavy metal ions (Pb²⁺) and triggering the CMU to express and release MtnB for Pb²⁺ chelation. **b)** A box-form MCU for individual mouse epidermal Pb²⁺ monitoring was designed to activate communication cross a mice population wearing CMU under the same environment potentially exposed to Pb^2+^. **c)** Characterization of mouse epidermal Pb^2+^ monitoring using a box-form MCU. **d)** Diagram of the integrated electronic circuit board in the wearable CMU. **e)** Photograph of mice wearing the CMU devices at epidermis. **f)** Fluorescence image of mice epidermis 5 hours after activating mCherry reporter secretion via the wearable CMU and its subsequent removal. **g)** Diagram of heme-sensing MCU triggering bacterial expression and release of IL-10 in CMU. **h)** Timeline of fecal heme monitoring in mice via box-form MCU and the activation of intervention through intestinal expression and release of IL-10 controlled by wearable CMU. **i)** Continuous monitoring of fecal heme levels in different mouse groups over 3– 6 days using the box-form MCU. **j)** Daily body weight changes of mice in different experimental groups. **k)** Comparison of colon lengths among groups at the end of the experiment. **l, m, n)** Levels of inflammatory cytokines (TNF-α, IL-12, IL-1β) in colon tissues of different groups at the end of the experiment.

To evaluate the availability of MCI-based communication for long-term inflammation monitoring and intervention, we established mouse model of inflammatory bowel disease (IBD). The MCU-based biosensing of heme, a biomarker associated with colitis, in fecal samples would trigger the CMU-activated expression and secretion of anti-inflammatory cytokine IL-10 (Fig. 5g). A monitoring-driven intervention model for mice colitis was developed. Inflammation was induced via oral administration of 2% DSS for the first 6 days, and based on the MCU monitoring results, the CMU was activated to implement therapeutic intervention (Fig. 5h). Firstly, we tested the response to different heme concentrations *in vitro* and established a threshold of 10 μA, corresponding to heme levels above 0.05 μM, as an indicator of severe inflammation (Fig. S10a, S10b). We then monitored fecal heme levels using the MCU on days 3, 4, 5, and 6 after DSS treatment (Fig. 5i, S10c, S10d). On day 6, the readings exceeded the 10 μA threshold, indicating significant colitis. Subsequently, light-responsive EcN engineered to secrete IL-10 were introduced into the diet of the treatment group, and abdominal green light irradiation via wearable CMU (Fig. 5d) was applied for 3 hours daily. Results showed that the weight loss trend in CMU-treated mice gradually slowed, stabilizing above 80% of initial body weight, better than other DSS-treated groups (Fig. 5j). When the mice were killed on day 12, colons were collected and the lengths were measured. CMU-treated mice exhibited noticeably longer colons compared to those of other inflamed groups (Fig. 5k, S10e). Analysis of typical pro-inflammatory cytokines revealed that levels of TNF-α, IL-12, and IL-1β in CMU-treated mice were significantly lower than those in other DSS-treated groups and closer to those of healthy control mice (Fig. 5l–n). In our previous work, we also demonstrated that a prototype of MCI consisting of an electronic capsule and engineered bacteria communities could be successfully used for early diagnosis and intervention of pig colitis^8^. In summary, these results demonstrated the applicability of MCI-based communication for long-term monitoring and intervention in chronic inflammatory bowel disease.

### Program MCI-based communication for unmanned surface vessel-based aqueous monitoring

Besides biomedical applications, we conducted a bold test to explore whether MCI-based communication could drive mechanical functions. For this purpose, we established a model for detecting As³⁺/AsO_2_^−^ in an environmental monitoring scenario, which drove the mechanical functions of an unmanned surface vessel (USV) (Fig. 6a, S11a). The MCU and auxiliary devices in the USV were capable of collecting water samples from a designated water area (Fig. 6b). When the MCU-detected As³⁺ in the water exceeded a threshold concentration, it would trigger the turning on of an LED indicator on the vessel as an alarm function (Fig. 6b). Firstly, we verified that a basic MCU box model containing As³⁺/AsO_2_^−^-sensing bacteria^45^ exhibited a positive concentration-dependent response to NaAsO_2_ solutions at concentrations of 0, 0.125, 0.25, 0.5, and 1 μM (Fig. S11b, 6c). Subsequently, we designed a systematic program in APP to control the entire process of aqueous sampling and sensing (Fig. S11c). This process was operated by an embodiment of MCU device installed within the hull of the USV (Fig. 6d, S11d). The MCU device was connected to the hull via a conduit controlling the “water switch” to collect samples from the environmental water body. Once the water sample entered the “liquid storage” chamber, it was mixed with the As³⁺/AsO_2_^−^-sensing bacteria and stirred to accelerate the biosensing process. A round electronic circuit board was used to convert the bioluminescence output of the bacteria into photocurrent signals. All the mentioned functions above were managed by a rectangular electronic circuit board as “control module” (Fig. S11d).

**Fig. 6.**
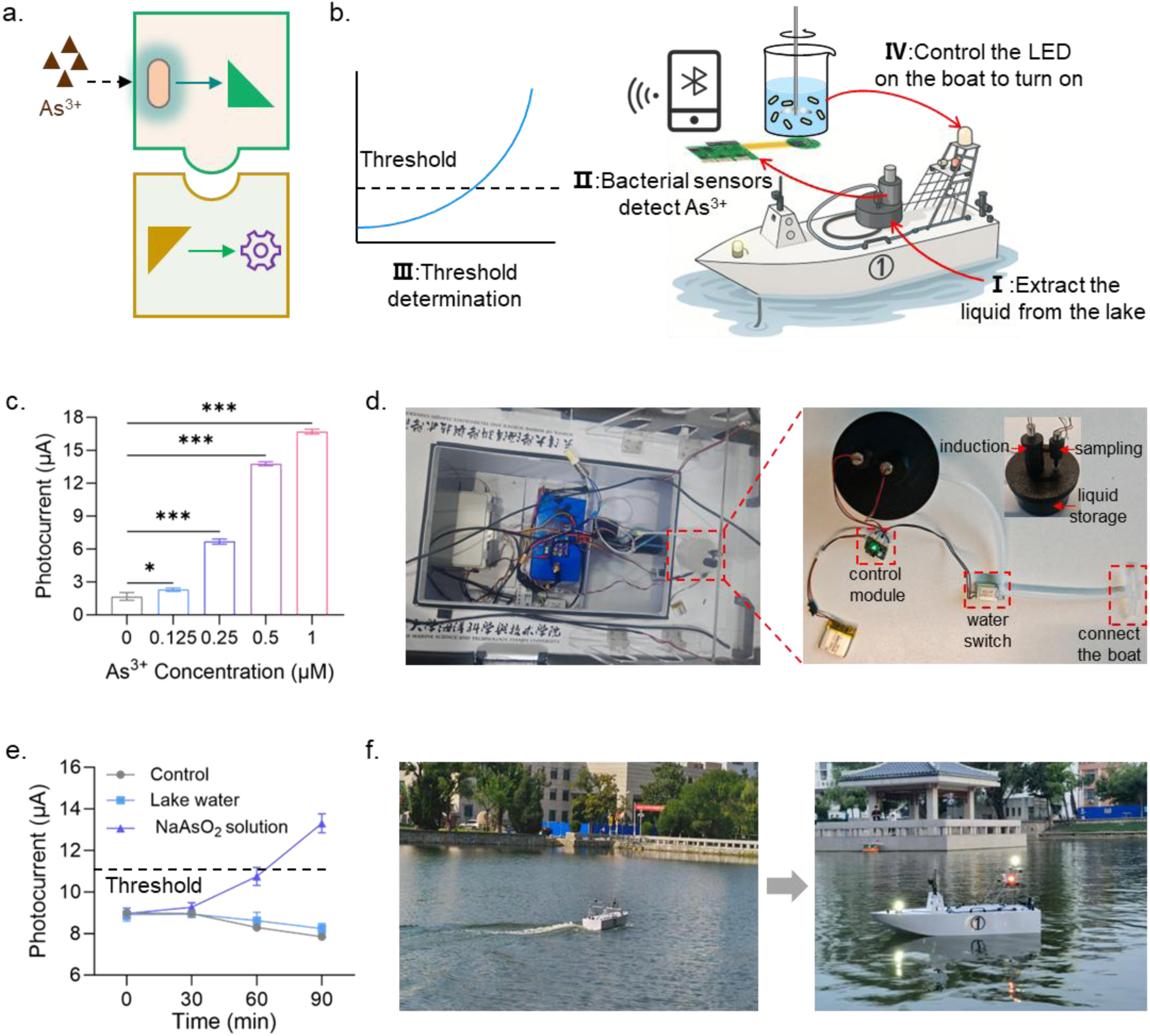
Evaluation of MCI-based communication for unmanned surface vessel-based aqueous monitoring. **a)** Diagram of the MCU-based As³⁺/AsO₂⁻ sensing and subsequent communication-triggered mechanical actuation. **b)** Workflow of water sampling by an unmanned surface vehicle (USV), MCU-based detection of As³⁺/AsO₂⁻ exceeding a predefined threshold, and activation of an alert LED on the USV. **c)** Detection of different concentrations of AsO₂⁻ using a box-form MCU model. **d)** Photograph of the USV and the MC-based communication system installed in its hull compartment. **e)** Detection results of lake water samples spiked with NaAsO₂ using the system shown in (d). **f)** Photograph demonstrating the USV operation of sampling in a specific water area, sensing AsO₂⁻ beyond the thershold and activating the alert LED.

Sampling test results demonstrated that saturated volume sampling in the MCU could be achieved within approximately 3 seconds (Fig. S11e). Finally, under laboratory conditions, we used the MCU to test real lake water samples collected from Jing Ye Lake at Tianjin University, as well as lake water samples with NaAsO₂ addition to get 1 μM concentration of AsO₂^−^. The results showed that positive detection of AsO₂^−^ could be achieved within just 90 minutes (Fig. 6e, S11f, S11g). Moreover, this detection outcome could also be used to control the switching on of an LED alarm light on the USV (Fig. 6f). In this final scenario, the laboratory-prepared water samples containing NaAsO₂ were pre-loaded into the MCU device, rather than being collected directly from the open lake environment. To summarize, we successfully constructed a system that could be used for life-centered biosensing of special chemical in environmental samples and beyond information exchange processes, which could be further connected to the IoT.

## Discussion

This study aims to extend the human-computer interaction (HCI) successful experience to microbe-computer interaction (MCI), a representative living biosystem-computer interaction (LBCI). Our goal is to establish life-nonlife and cross-life communication among engineered microbial communities. The main challenge of MCI is how to convert state-related signals from noncognitive microbes into quantifiably measurable digital signals, and conversely, how to achieve programmable and quantitative regulation of biosystem states via computer means. To address this, we first established two basic opto-interfaced MCI units, namely the microbe-to-computer unit (MCU) and the computer-to-microbe unit (CMU). In the MCU model, we demonstrated a strong positive correlation between luminescent bacteria density (10^4^ to 10^8^) and detected photocurrent values (0 to 32.5 μA), realizing at least 100 min successive monitoring of bacterial luciferase expression. We also verified that the approaching of bacterial cells to electronics would dynamically activate the working of MCU, allowing customized MCU design. In the CMU model, we thoroughly evaluated a rechargeable CMU pattern for activating optogenetically engineered microbial functions as far as 19 cm away.

In addition, we developed devices of MCU sensors, CMU actuators, and transceivers integrated by MCU and CMU, for biomolecule information processing. The MCU sensor enabled quantitative sensing of various biomolecules, such as nitrate, tetracycline, heme, oxygen, Pb^2+^, As^3+^/AsO2^−^. It was very exciting to expand the MCU function for sensing large biomolecules such as DNA^46^ in the future. The CMU actuator, in turn, allowed precisely spatiotemporal control of bacterial cells to produce and release functional biomolecules. The microbe-centric transceiver enabled both specific cell stimulation and outcome monitoring, establishing feedback with computers even within a complex live pig colon. Thus, we have developed a wide range of customized units capable of fulfilling the fundamental functions of MCI. The modularity, reusability, deployability and accuracy of our MCI units were fully validated.

Building upon the fundamental units, we further established diverse MCI-based communication architectures using programmable strategies. Our investigation focused on the remote point-to-point effectiveness, logic architecture programmability, and temporal tunability of the communication system. The effectiveness of point-to-point communication was validated by the successful connection of unicast and multicast modes. The programmability of logic architectures was demonstrated using both an AND-gate logic and a multi-level complex logic with three inputs and three outputs. Furthermore, the temporal tunability of the system was verified via cascaded and dynamically regulated feedback modes. It is supposed feasible to realize more customized cross-life communication architectures, which do not have to rely on close contact between cell populations and common nutritional conditions^47^.

Our designer MCI-based communication demonstrated verifiable value for biomedical applications. In the tumor scenario, we validated a wearable unicast mode of MCI-based communication to monitor the tumor hypoxic microenvironment and drive the control of bio-therapeutic intervention. In the epidermal scenario, we demonstrated that a multicast mode of MCI-based communication achieved early warning of individual skin status within a population under prolonged environmental exposure to heavy metal ions, triggering a collective response to chelate these ions and maintain skin health. In the intestinal scenario, we confirmed that MCI-based communication featuring time-dynamic threshold response characteristics could be applied to long-term monitoring and therapeutic intervention in chronic colitis. Our strategy is promising for broad biomedical scenarios such as in long-term chronic diseases, conditions prone to metastasis, and systemic disorders.

Finally, we established a cross-domain communication mode linking biological and mechanical systems, integrating biological monitoring of heavy metal contamination in water samples with the functionality of an unmanned vehicle. Our MCI-based cross-domain communication presents a significant advantage over conventional water extraction and detection protocols by enabling real-time, dynamic monitoring. Leveraging the mobility and cruising functions of unmanned surface vessel (USV), the integrated system allows for precisely targeted and real-time surveillance of specific zones in a water body. Moreover, it holds promise for not only detecting concentrations of toxic and harmful substances but also for tracing potential release origins by following concentration gradients. The expanded unmanned vehicles such as surface vessels, drones, and ground robots would enable targeted monitoring of environmental biomarkers^48–51^, serving as long-term biohybrid intelligent monitoring platforms.

This study can be further extended to contribute to multiple research and application fields. For scientific research, our programmable MCI-based communication architecture provides a platform that supports extended and sophisticated connection patterns among different biological communities with the assistance of local area network (LAN) and internet. This will expand the scope of symbiotic systems and redefine approaches to artificial biosystem design^52,53^. Besides, the MCI and extensive LBCI hold transformative potential for complementing internet of things (IoT) with biosystems, promoting the internet of things and biosystems (IoTB)^10,54^. For instance, IoTB can be envisioned as networked biohybrid cyborg systems or ecosystems, potentially expanding the boundaries of human perception and agency. For applications, the iterative development of smart wearable, ingestible, implantable, unmanned vehicle-carried and more imaginative forms of MCI and LBCI are especially valuable for connecting diverse life activities, life-environment and multi-life interactions to IoTB. We can establish statistically valuable tools for social health monitoring and maintenance, and ecological analysis and protection across higher dimensions.

This study is still at a preliminary stage, with several limitations and challenges remaining to be addressed. There is significant room for improvement. On the biological side, key issues that require further resolution include the maintenance and renewal of living cells, as well as the diversification^30^ and enhanced accuracy of quantitative synthetic biological functionalities^55^. Incorporating internal division of labor within cellular communities may also enable more sophisticated functions^32,33,56^. On the computational side, existing integrated electronics shall be further optimized in terms of intelligence, miniaturization, and energy efficiency^13,57^. The reliable integration of biosystems with highly integrated electronics will facilitate monitoring of non-deterministic biological functional variations^58^. For cross-life communication, further refinements in programming are necessary, including biohybrid reset mechanisms, improvements in speed and efficiency, and the AI-driven biological directed evolution^59,60^ to enable training of biohybrid intelligence.

## Methods

### Microbes and culture conditions

The strain *E. coli* DH5α and *E. coli* Nissle 1917 (EcN) were used as the chassis for the construction of the major engineered strains used in study. Luria-Bertani (LB) broth and plates were used for strain selection and cultivation. Kanamycin (50 mg L^−1^), ampicillin (100 mg L^−1^), and chloramphenicol (50 mg L^−1^) was added appropriately according to different antibiotic resistance conditions. The strain *B. subtilis* 168 (ATCC 23857) was used in study and the cultivation medium was also Luria-Bertani (LB) added with corresponding antibiotics for strain selection. The yeast *S. cerevisiae* BY4741 used in this study and the cultivation medium was Synthetic complete (SC) medium (0.67 % yeast nitrogen base without amino acids, 2 % glucose, and appropriate amino acid drop-out mix), lacking certain amino acid for selection of yeast transformants.

### Microbial genetic engineering

All the microbial plasmids and microbial strains used in this study were listed in Table S1 and S2. The strains expressing luciferase in a constitutive manner included *E. coli* DH5α-Lux, *B. subtilis*-Lux and *S. cerevisiae* BY4741-Lux. The strains DH5α-Ara, EcN-Nitrate, EcN-Tet, EcN-Heme, DH5α-Hypoxia, DH5α-Pb, EcN-As were respectively constructed as biosensor strains for sensing L-arabinose, nitrate, tetracycline, heme, hypoxic condition, Pb^2+^, AsO_2_^−^. For the construction of green-light-responsive strains, an optimized CcaS-CcaR two-component optogenetic circuit was used to activate certain reporter protein expression. The reporter proteins included GFP and secreted GFP, bacterial luciferase expressed by *luxA* along with *luxBCDE* and other functional secreted proteins including PD-L1nb, MtnB, IL-10. The expression of α-amylase in yeast was controlled by a blue-light-responsive optogenetic circuit. When the reporter protein was aimed to be released from EcN, a type-I secretion system including HlyA peptide and assisting HlyBD proteins were introduced.

### Smartphone APP program and data processing

For making the working of BCI units and BCI-based communication under human supervision, an APP program on smartphone was developed. The diagram of connecting the varying CMU forms with Bluetooth through APP program was shown in Fig. S1b. The basic analog-to-digital converter (ADC) data detection and recording interface was shown in Fig. S1c-e, presenting the detection of different luminescence sources. The button of “Record” in the interface was clicked to upload the test data to the local file of the mobile phone. Besides, special operations could be programed in this APP interface according to customized demands in varying scenarios showcased in the Fig. 3f and S11c.

For data processing on computer, the form of the numerical value (ADC) recorded by the analog digital converter could be converted into a direct reflection of the photoelectric signal, photocurrent. After converting the ADC to a photocurrent, the average of the photocurrent during the time period of the collected data was calculated and used as the final photocurrent value of certain concentration of bacterial liquid. A linear relationship between the bioluminescence intensity and the photocurrent was fitted and obtained by drawing a picture with the abscissa for different chemiluminescence intensity and the ordinate for the scatter plot of the corresponding optical current values. This ADC-photocurrent formula was also used in all the different MCU forms in this study.

### Detection of microbial bioluminescence with various MCU patterns

For testing the bioluminescence of microbial constitutively expressed luciferase, 1% of the DH5α-Lux cultured overnight under uninduced conditions were inoculated into a fresh cultivation medium. Then, the appropriate densities of strains in logarithmic growth phase were measured by microplate reader or the primary CMU model. Detailly, the electronic circuit board was attached to the bottom of an opaque carton with adhesive tape, with the photosensor side was faced up to keep the circuit level. A square cover glass flat was placed above the photosensor, and the DH5α-Lux liquid medium was added on the cover glass with a pipette droplet to cover the square of photosensor for the bioluminescence measurement. This measured value of the photosensor was the recorded. The overall measurement by CMU shall be operated under a completely dark environment. Similar to this testing method, the bacterial liquid diluted in a tenfold gradient was detected. The OD_600_ values of each bacterial liquid sample was measured separately, where the OD_600_= 1 was approximately equal to 10^9^ CFU/mL. When the optical signal of the sample to be measured was nearly equal to the optical signal value of the air, it was regarded as the limit for detecting low density of strains.

#### Capsule-form MCU test

The above integrated electronic circuit board in MCU was retained, except that its battery was changed into three commercial button batteries (80 mAh, 393/LR754, KOONENDA) in series by welding on the board. The whole board was assembled with two halves of PET shell with the UV curing adhesive evenly applied at the connection and cured through the UV lamp irradiation to realize the unleaked packaging of capsule. The ability of electronic capsule for monitoring the bioluminescence of approaching DH5α-Lux *in vitro* was tested by using a 30 cm intestinal segment. 2 mL of OD_600_=1 DH5α-Lux was injected at a specific position in the intestinal segment. Then, the electronic capsule was placed from one side of the intestinal segment and be pulled by a constant speed to the other side to simulate the downward movement of the capsule in the intestine. The changing data of electrical signals in the electronic capsule over time was recorded.

#### Microneedle-sampling-form MCU test

The structure of the whole MCU was divided into three layers, namely, the top layer (containing a half pressing chamber layer and a half circuit layer), the middle layer (containing a half cavity layer and a half sensor-bacteria curing layer), and the microneedle layer at the bottom. The two halves of the top layer were connected together by PDMS reverse mode. In order to not disturb the bioluminescence detection of bacteria, the top pressing layer and middle cavity layer was connected by edible silica gel. The 2% sodium alginate was mixed with the bacterial solution in a 1:1 volume and immersed in CaCl_2_ solution for curing. The cured bacteria were placed in the middle layer adjacent to the cavity layer. The middle cavity layer and the bottom microneedle layer were stuck by the appropriate size design. For bacteria sampling, the DH5α-Lux was placed in liquid culture dish, and this microneedle-sampling-form MCU was pressed to absorb the bacterial samples to test its bioluminescence performance.

#### MCU sensor test

The respective biomolecule samples of L-arabinose, nitrate, tetracycline, heme, Pb^2+^, AsO_2_^−^ in this study were mainly detected by the MCU sensor. A light-proof box was fabricated by 3D printing, composed of two sections (as shown in Fig. S4b). The engineered bacteria biosensors were housed in the two wells of the lower section, while the integrated circuit board was immobilized in the upper section. Biomolecule samples were introduced into either well, and then the upper section was slid to align with this well, enabling photocurrent measurement. The testing procedure for each biomolecule of concentration series was as follows: first, the appropriate volume of high concentration biomolecules was added to the OD_600_=0.2 culture tube to form the concentration series to be tested; then, each tube was incubated at 37 ℃ shaker for 2 hours; finally, the tube containing 2 mL samples was inserted into the well in the box-form MCU sensor for photocurrent measurement. For the murine blood spiking experiment, the 1 to 2 mm of the mouse tail was cut out and a small amount of blood was removed from the tail vein. Then, the blood sample was firstly transferred to a tube with anticoagulant and appropriate amount of tetracycline, then directly added to the tube containing biosensor strains. After 2 h cultivation at 37 ℃, the mixture tube was measured by MCU sensor.

### Activation of optogenetically engineered microbes in CMU actuator

The stent-form CMU actuator was composed of a stent, electronic circuit, optogenetically engineered microbes and the supporting material. The stent used to support the MCI was purchased from Nanjing Minimally Invasive Co., LTD. with the size of 26 mm width and 80 mm length. Optogenetically engineered microbes were immobilized on the scaffold by forming a form of complex with PVA. The method of preparing PVA-microbe mixture was as follows. Firstly, a 10 wt% PVA solution was prepared by dissolving 10 g PVA (molecular weight 146000-186000, 99+%; Bid Pharmaceutical) in 90 ml of 90 ℃ water for 5 h until clarification. Later, PVA was mixed with the bacterial solution, applied on the stent, standing for 1 hour at 37 ℃, and the curing was checked.

### Mouse experiments

Male C57BL/6 mice (7∼8 weeks) were purchased from SPF (Beijing) Biotechnology Co., Ltd. Female Balb/c mice (6-8 weeks) were purchased from GemPharmatech Co., Ltd. (Jiangsu, China). Mice are raised in clean cages with a constant temperature of 20°C and the light was kept on from 8 a.m. to 8 p.m. every day. Mice are fed with sterilized water and mouse maintenance feed (SPF-F02-001, SPF (Beijing) Biotechnology Co., Ltd., China), and their cages are kept clean and comfortable with shavings padding (SPF-M01-001, SPF (Beijing) Biotechnology Co., Ltd., China). Experiments were carried out after the mice were reared adaptively for one week.

#### Tumor biomarking and intervention scenario

4T1 breast cancer cells in the logarithmic growth phase were harvested and resuspended in serum-free medium. Each Balb/c mouse was inoculated with 1×10⁷ cells in a 200 μL volume via subcutaneous injection. The tumor model was considered established 10 days post-inoculation. The hypoxia-sensing strain DH5α-Hypoxia from an overnight culture were intratumorally injected at concentration of 2 ×10^9^ CFU/mL in a 50 μL injection volume. Then, the integrated circuit boards combining both CMU and MCU electronic circuits were worn respectively on these mice. Bioluminescence imaging on the mice was performed using an IVIS Lumina S5 system at various time points post-injection (0 h, 1 h, 2 h, 4 h, and 6 h). Photocurrent measurement on the mice was conducted 2 hours after the intratumoral injection of DH5α-Hypoxia and inside a light-tight chamber. Photocurrent was recorded for 2 minutes per mouse at 0 h, 1 h, 2 h, 4 h, and 6 h post-injection.

#### Epidermal Pb^2+^ monitoring and chelation scenario

Firstly, the Pb^2+^ on C57BL/6 mice skin was monitored using MCU sensor. Overnight culture of the Pb²⁺-sensing bacterial strain was subcultured into fresh medium at a 1:50 ratio and grown for 6–8 hours until OD₆₀₀ reached 0.5. The bacterial cells were harvested by centrifugation at 4,000 × g for 15 minutes, and the pellet was retained. The dorsal skin of the mouse was shaved to expose a defined area. A 100 μL solution of 10 μmol/L Pb²⁺ was applied to the exposed skin and allowed to air-dry. The Pb²⁺-coated area was then rinsed with 1 mL of LB medium via gentle pipetting to collect the sample, and the volume was adjusted to 1 mL to form the sampling solution. This 1 mL sampling solution was used to resuspend the Pb²⁺-sensing bacterial pellet. The resuspended culture was incubated at 37°C with shaking at 220 rpm for 2 hours. Bioluminescence was subsequently measured using a microplate reader. In parallel, the bacterial suspension was also analyzed using the box-form MCU sensor. Data were recorded via a dedicated mobile app over a 2-minute interval and processed to obtain photocurrent values.

Secondly, the release of target protein from a Skin-Integrated CMU Actuator in Mice was triggered by the MCU sensor result. The engineered strain EcN-Opto-secCherry, designed for optogenetically controlled mCherry expression and secretion, was grown overnight under light-shielding conditions. The culture was centrifuged at 4,000 × g for 15 minutes to separate the supernatant, and the bacterial pellet was retained. A total of 1 g of chitosan–β-glycerophosphate hydrogel was thoroughly mixed with the bacterial pellet. The mixture was loaded into annular patches, sealed with a PU membrane to form hydrogel dressings containing the optogenetically controlled bacteria. The electronic module was then affixed over the PU membrane to ensure a light-protected environment. Based on a predefined program in the smartphone APP, LED irradiation was activated on the mouse skin when the MCU sensor detected Pb²⁺ levels exceeding a preset threshold. After several hours of irradiation at room temperature, the hydrogel solidified into a film. The activation of mCherry expression was assessed using *in vivo* imaging. Following removal of the CMU actuator and hydrogel film, a subsequent *in vivo* imaging session was performed to evaluate the retention of bacterially secreted mCherry on the skin surface.

#### Colitis diagnosis and intervention scenario

After one week of adaptive feeding, the C57BL/6 mice were randomly divided into four groups: Control, DSS-induced colitis model group (DSS), DSS-induced colitis with subsequent light intervention (Light + DSS), and DSS-induced colitis without subsequent light intervention (Dark + DSS). Before the whole experiment, body weight was measured for all mice, and a 2% (w/v) DSS solution was prepared in pure water and administered to all DSS-modeled groups via drinking water, while their diet remained normal; the Control group received only regular drinking water and food. After 6 days of 2% DSS administration, the DSS-modeled mice were switched back to pure water. Body weight of each mouse was recorded daily from Day 1. Starting from Day 3, fresh fecal samples were collected daily and directly introduced into liquid culture medium containing the heme-sensing bacteria. Specifically, an overnight bacterial culture was subcultured and grown to OD₆₀₀ = 0.5. After adding mouse feces, the mixture was incubated for 2 hours, followed by bioluminescence imaging or MCU sensor detection. A photocurrent value exceeding a predetermined threshold on Day 6 indicated severe intestinal bleeding. From Day 7 onward, the Light + DSS group of mice received oral gavage of EcN-Opto-secIL-10 bacteria that had been pre-induced overnight with green light, and wore a CMU actuator to provide 5 hours of daily light exposure. In contrast, the Dark + DSS group of mice was gavaged with the same bacterial strain but without any prior light induction. After the experiment concluded on Day 12, all mice were euthanized, and colon tissues were collected for length measurement. Levels of pro-inflammatory cytokines (TNF-α, IL-12, IL-1β) in colon tissue were analyzed, and fecal calprotectin (CalP) concentration was measured by ELISA.

### Pig experiments

Quarantine healthy ordinary domestic pigs weighing 20-25 kg were selected for a week cultivation before animal experiment. Pigs were fasted for foods for 24 h and water for 4 h and operated by coloclysis using lactulose for 2 h before surgery. After emptying the intestine, the pig was anesthetized by intramuscular injection with zololeteline (5 mg/kg) + cycerazine hydrochloride (2 mg/kg). The EcN strain containing the genetic circuit of green-light-responsive luciferase expression (EcN-Opto-Lux) was encapsulated in the stent-from CMU actuator. This CMU actuator was implanted into the designated position of the pig rectum with the assistance of an implantation tool. The tool had a diameter of 8 mm and a rubber pad on its head in order to push out the stent-form CMU smoothly. An animal X-ray imaging system (Nanjing Puai Medical, China) could take photograph to determine the implantation position. In order to verify whether the implantation of stent-form CMU would produce inflammation, serum CPR and SAA levels (reflecting potential inflammation or injury) before or after scaffold implantation were tested. After the experiment, 10% KCl solution (0.5 mL/kg) was quickly injected to kill the pigs, the colonic tissue was sampled for hematoxylin and eosin (H&E) staining and analysis.

Test of the biohybrid transceiver in live pig intestine was operated as follows. After the implantation of stent-form CMU actuator via the tool, an endoscope (Xi’an Betnoch, China) was used to observe the LED switching in the stent-form CMU actuator activated by the TX charging adjusted outside pig body. After that, the capsule-form MCU constructed above in this study was loaded transanually to the stent implantation position. Then, the capsule would measure the bioluminescence emitted from the EcN-Opto-Lux in the stent-form CMU actuator. The measured data was recorded by smartphone APP.

### MCI-based communication for aqueous sampling and monitoring by USV

The USV was constructed from high-strength aluminum-magnesium alloy, combining excellent corrosion resistance with structural strength for long-term marine operations. The upper structure featured a modular bracket design that allowed flexible installation of various mission-carrying payloads and control systems. The sampling-detection MCI platform integrated a custom-designed hybrid PCB circuit board with a 3D-printed sampling device (Tu Zhu H2D, Bambu Lab Co., Ltd.). A silicone hose connected the platform to the vessel’s liquid cooling circulation system, which branched into a sampling channel linked to an electromagnetic valve. This valve interfaced with the unmanned boat’s relay control module for real-time shut-on and shut-off, enabling preliminary water sampling. Powered by a brushed hollow cup motor, the 3D-printed centrifugal pump-inspired chamber facilitated sampling through silicone tubing and bacteria mixing induction. The integrated design enabled secondary sampling and monitoring of the bioluminescence of the bacteria sensor, achieving detection of target marker in water.

For simulation of aqueous sampling and monitoring, the 400 μL volume of pure water, lake water or lake water added with NaAsO2 was respectively mixed with 1.6 mL sensor strains pre-positioned in the sampling device of the MCI platform. After the bacteria mixing induction for 0 to 90 minutes, the respective photocurrents were measured. The result exceeding the preset value would drive the alarm LED to be turned on.

## Ethics statement

The study has been approved by local ethics review committee. In detail, all mouse procedures were performed in accordance with the Guidelines for Care and Use of Laboratory Animals of Tianjin University, following the requirements of the People’s Republic of China (GB14925-2010), and the experiments were approved by the Animal Ethics Committee of Tianjin University (TJUE-2021-009). All the pig procedures complied with the Guidelines for Care and Use of Laboratory Animals of Longyan University, following the requirements of the People’s Republic of China (GB14925–2010), and were approved by the Animal Ethics Committee of Longyan University (LY2024017X).

## Acknowledgments

This work is supported by the following fundings: National Natural Science Foundation of China for Excellent Young Scholars (32122047); Tianjin Natural Science Foundation for Distinguished Young Scholars (23JCJQJC00210); Beijing-Tianjin-Hebei Basic Research Cooperation Project of Beijing Natural Science Foundation (23JCZXJC00370); Key Program of Tianjin Natural Science Foundation (22JCZDJC00230); National Natural Science Foundation of China (82302366).

---

**Fig. S1.**
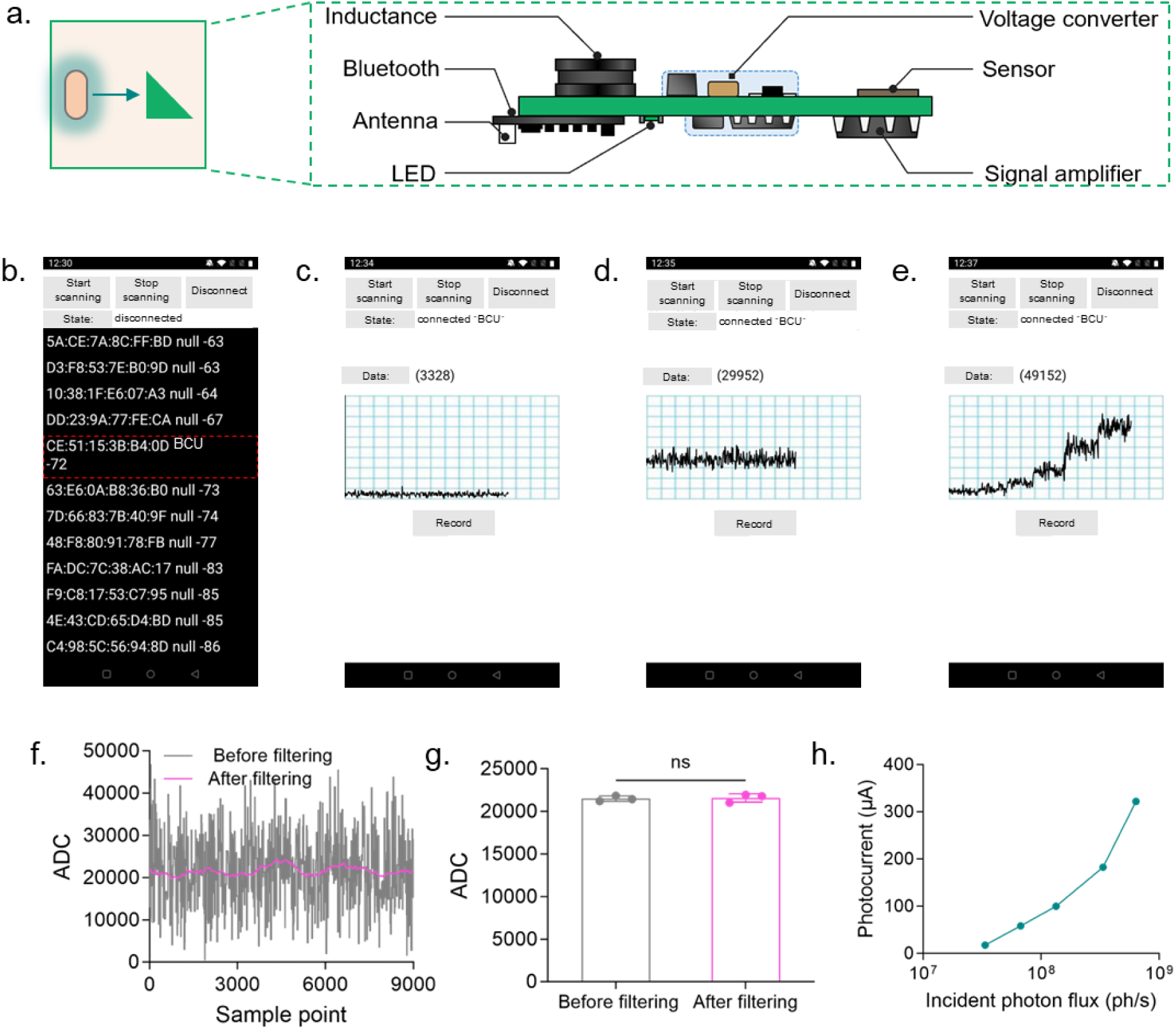
Diagram of MCU and photocurrent data recording and processing through smartphone APP. **a)** Diagram of the integrated electronic circuit board within the MCU, showing individual components. **b)** Smartphone APP interface showing Bluetooth scanning and connection to a specific MCU. **c)** Interface displaying electrical signal variation induced by background illumination under darkness. **d)** Electrical signal response under constant-intensity light under darkness. **e)** Electrical signal changes under progressively increasing light intensity under darkness. **f)** Application of a moving-average algorithm for real-time data filtering. **g)** The filtering process preserving the mean value of the 9000 data points collected over a 3-minute period. **h)** Increase of the detected photocurrent values with rising incident photon flux.

**Fig. S2.**
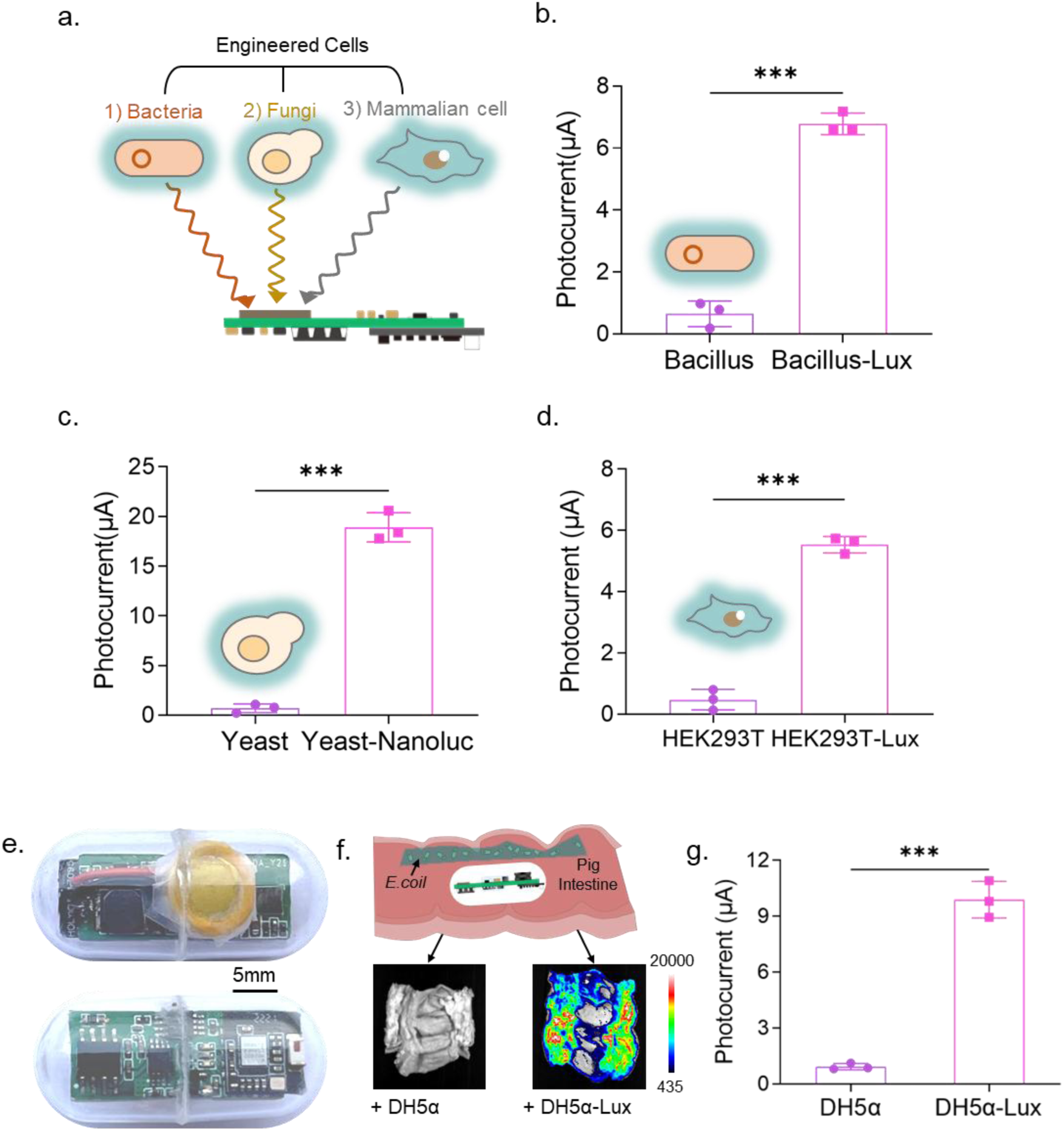
Expansion of MCU functions for detecting different specie cells or the approaching bacteria. **a)** Diagram of bioluminescence detection for bacteria, fungi, and mammalian cells based on the MCU model. **b, c, d)** Measurement of photocurrent values using the MCU for the respective bioluminescence of *B. subtilis*, yeast *S. cerevisiae*, and human HEK293T cells, respectively. **e)** Photograph of the integrated electronic circuit board assembled into an optically transparent capsule. **f)** Illustration of the capsule from (e) used for detecting either control DH5α bacteria or bioluminescent DH5α bacteria within a porcine intestinal segment *in vitro*. **g)** Experimental results of (f).

**Fig. S3.**
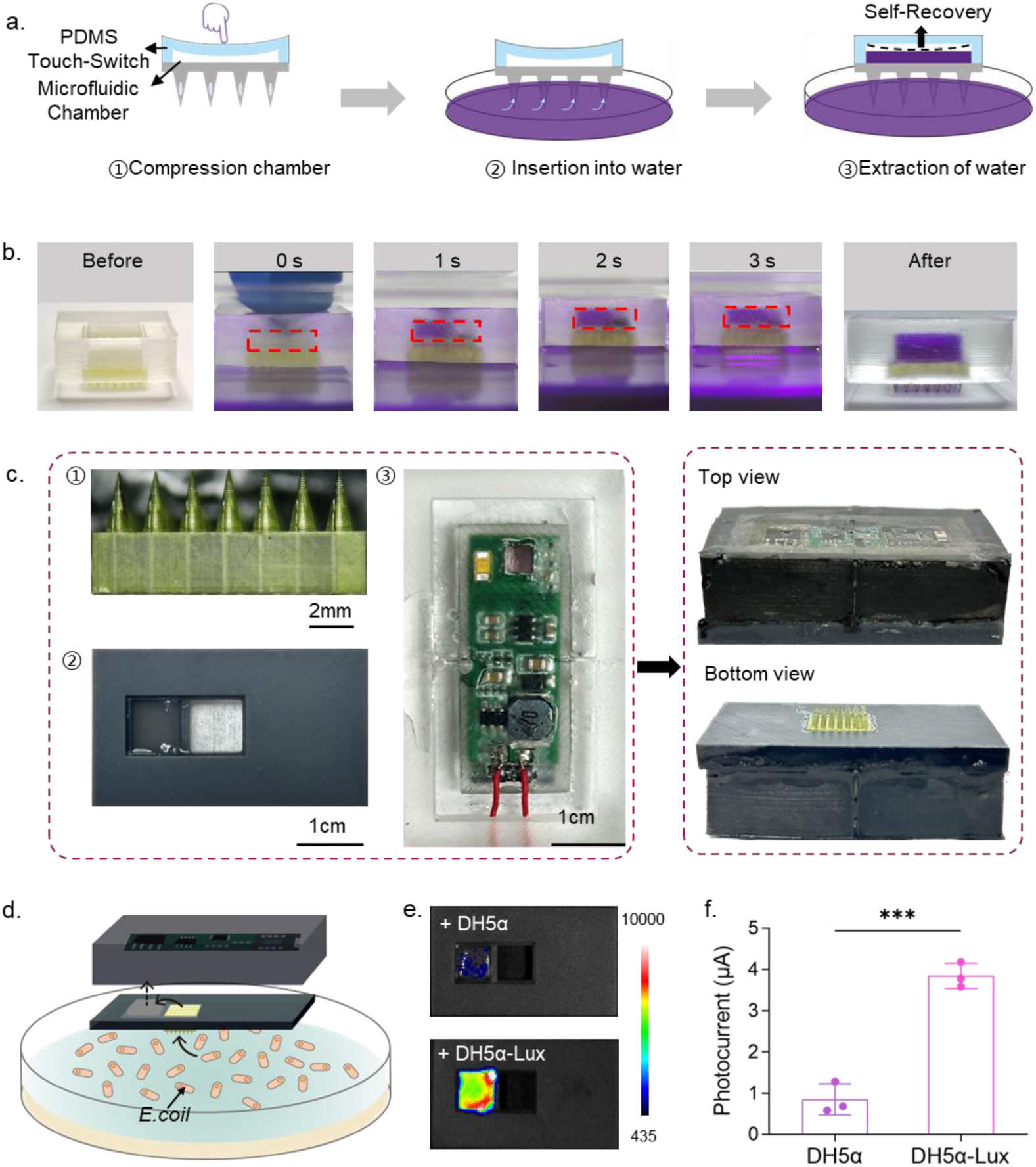
Expansion of MCU functions for detecting the sampled bioluminescent bacteria. **a)** Diagram of the sample uptake process using a touch-switch microfluidic chamber. **b)** Photographs of the chamber in (a) and the sample uptake process. **c)** Illustration of assembling MCU unit consisting of the microneedle, light-proof enclosure, and integrated electronic circuit board. **d)** Illustration of sampling from the liquid containing *E. coli* strains through the microneedle interface of the MCU unit. **e)** Imaging of bioluminescence after respective uptake of two different engineered strains using the setup in (d). **f)** Photocurrent detection results of the experiment in (d).

**Fig. S4.**
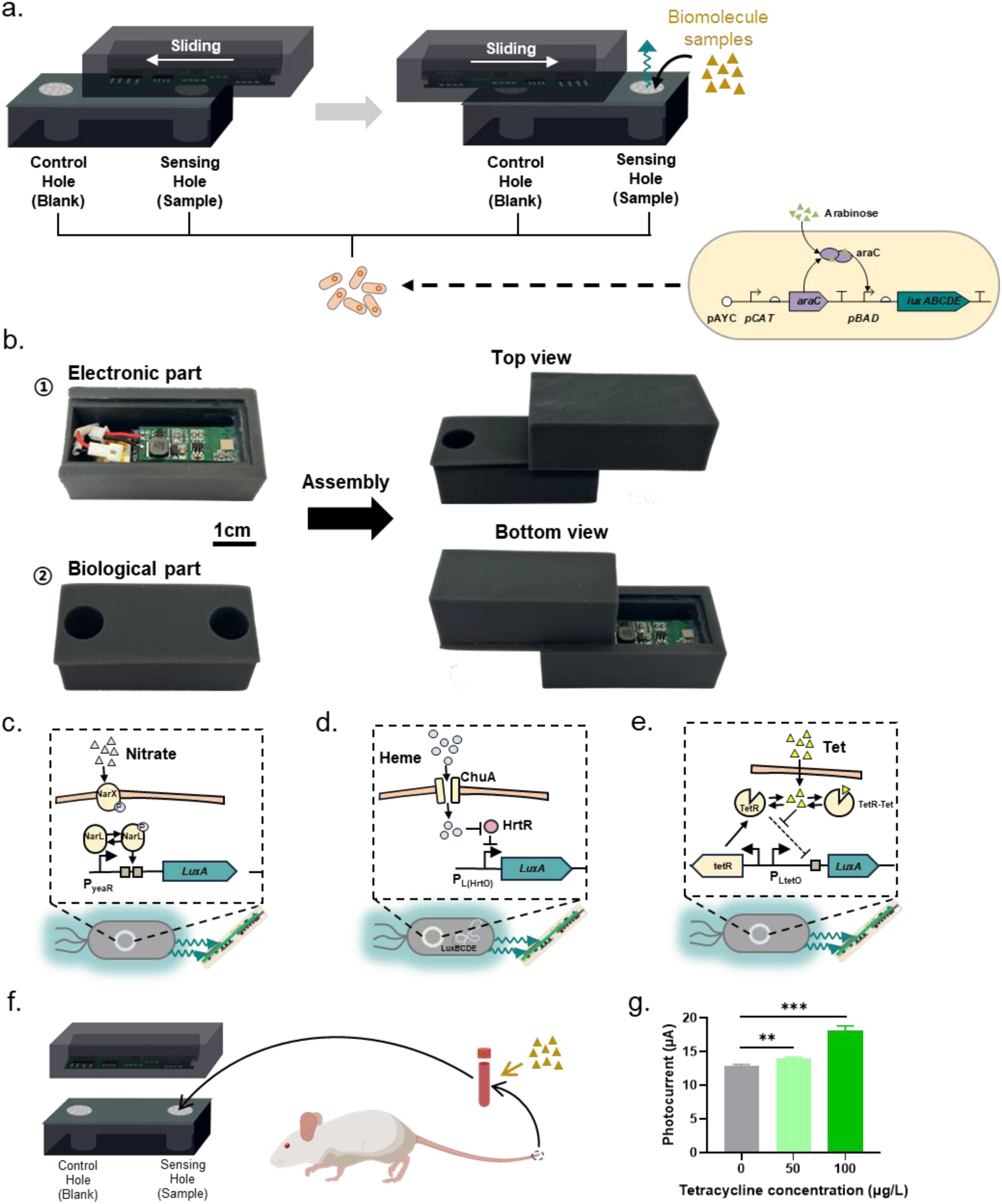
Evaluation of the box-form MCU sensor. **a)** Illustration of using the box-form MCU sensor for detecting additional biomolecule signals (e.g., L-arabinose). **b)** Photographs of the box-form MCU sensor shown in (a). **c, d, e)** Diagrams of the MCUs loaded with engineered bacteria carrying different genetic circuits for the detection of nitrate, heme, and tetracycline, respectively. **f)** Diagram of spiking experiment for detecting tetracycline in murine blood samples using the MCU sensor. **g)** Experimental results corresponding to the setup described in (f).

**Fig. S5.**
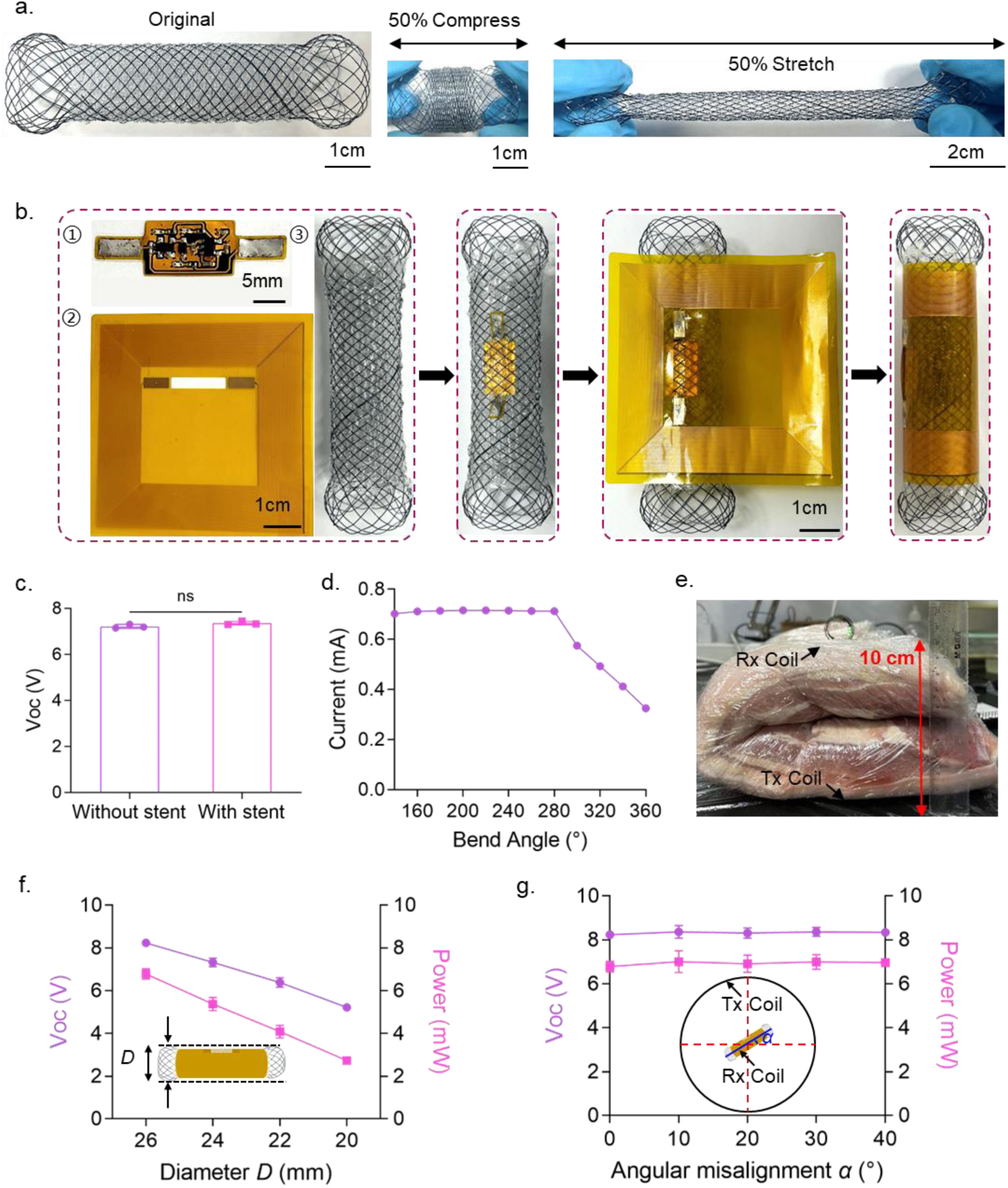
Assembly of the stent-form CMU actuator and electrical test. **a)** Feasibility test of 50% compression or 50% stretching of a stent encapsulated with biocompatible PVA material. **b)** Photograph of the fully assembled stent-configured MCU, consisting of the PVA layer (pre-loaded with optogenetic engineered bacteria as needed), stent structure, and electronic modules. **c)** Measurement of the open-circuit voltage of the receiving coil under different stent configurations. **d)** Effect of bending angle of the receiving coil on the operating current of the electronic circuit. **e)** Diagram of the experimental setup for wirelessly charging the MCU stent through a 10 cm-thick porcine tissue section using an external transmitter (Tx) coil and evaluating circuit performance. **f)** Test of the variations in open-circuit voltage and output power of the electronic circuit under stent compression. **g)** Test of the variations in open-circuit voltage and output power of the electronic circuit as the stent is rotated at different angles relative to the transmitter coil.

**Fig. S6.**
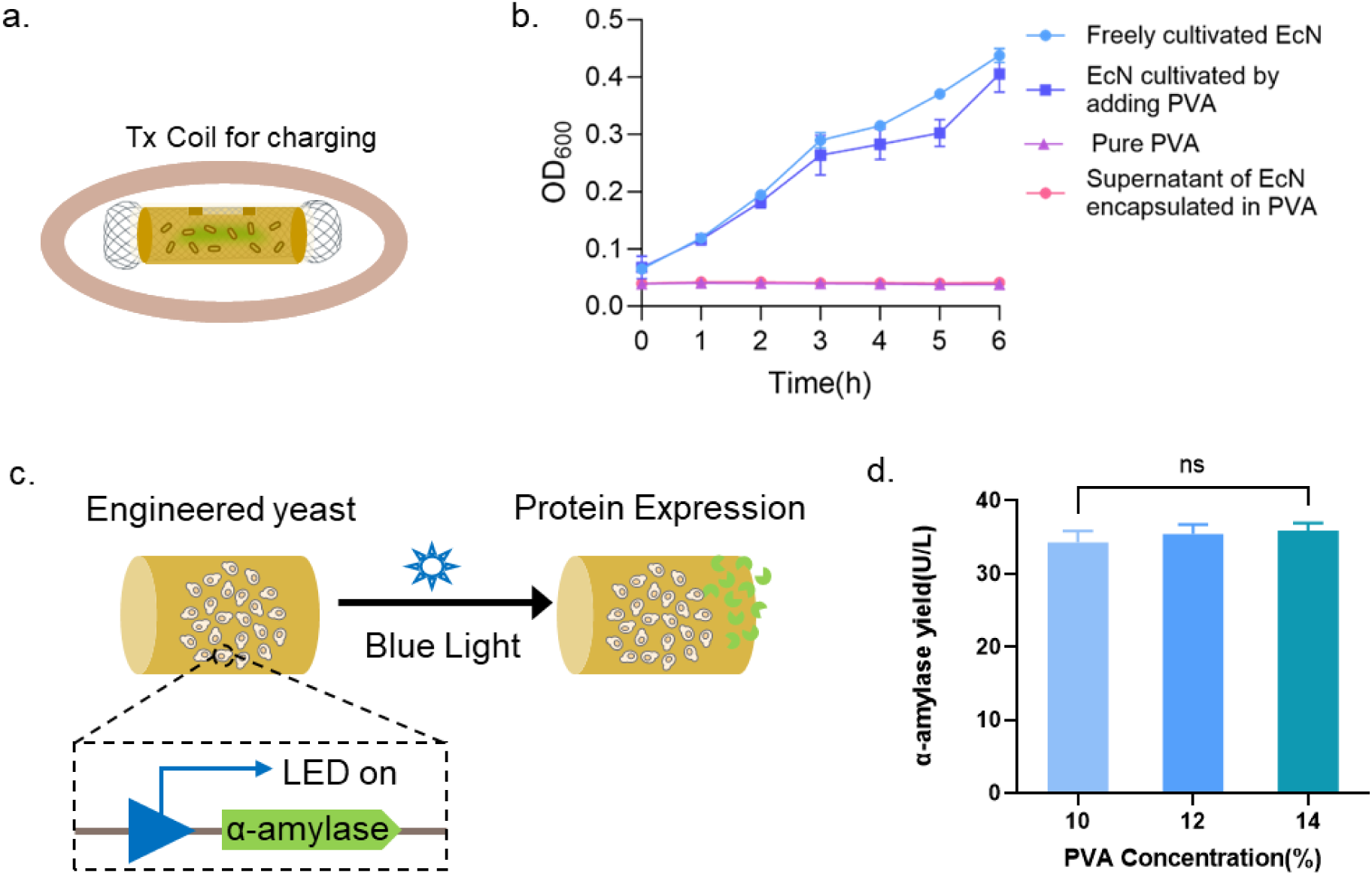
Test of opto-activation of microbial protein expression using stent-form CMU actuator. **a)** Diagram of wireless charging and activation of the stent-formed MCU actuator via a Tx coil. **b)** Test of the effect of PVA material on the growth performance of engineered EcN bacteria in culture medium, along with the assessment of potential bacterial leakage from PVA-encapsulated EcN. **c)** Diagram of the activation of α-amylase expression in PVA-encapsulated engineered yeast upon LED switching. **d)** Effect of varying PVA concentrations on the expression and secretion of α-amylase in encapsulated yeast.

**Fig. S7.**
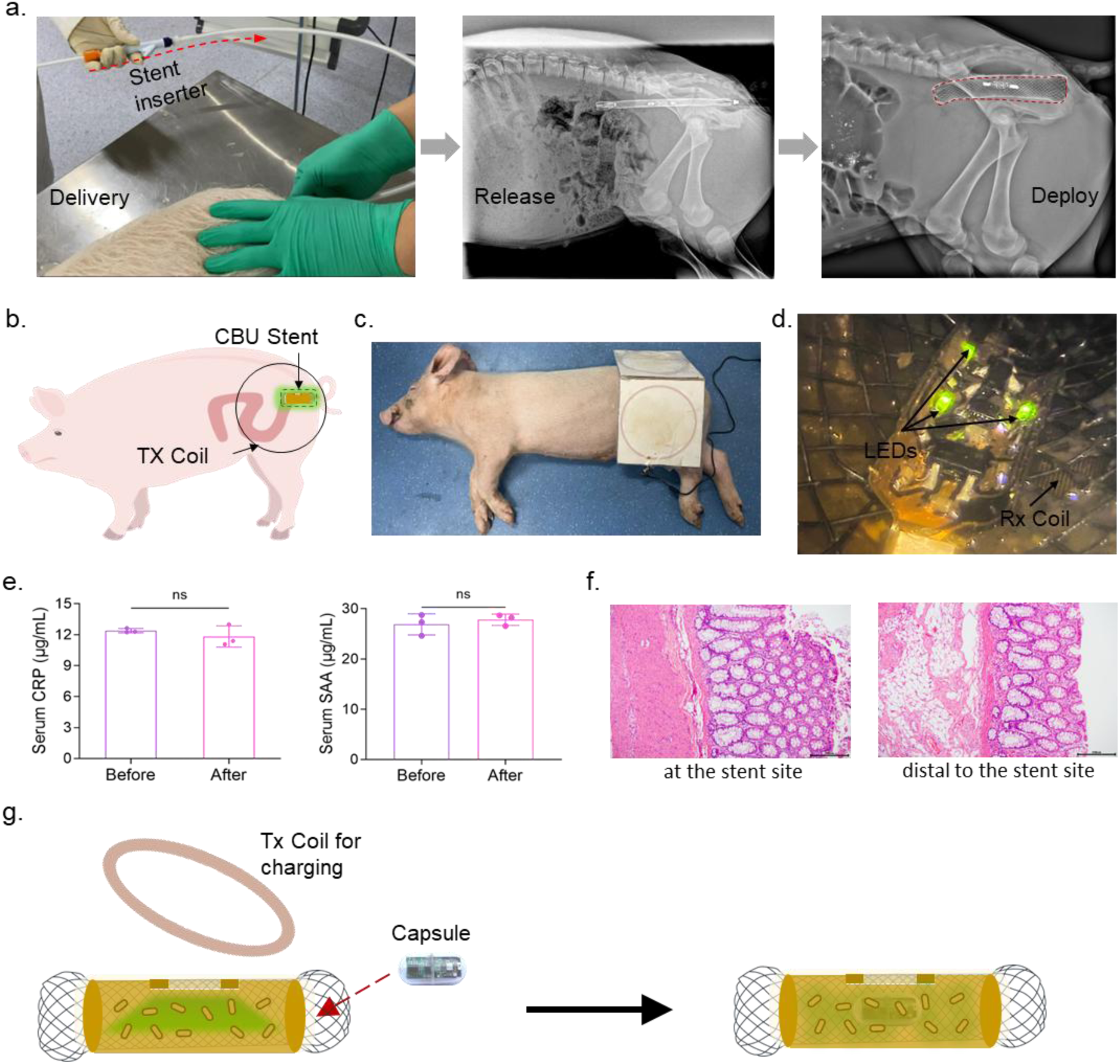
Evaluation of transceiver combined by CMU stent and MCU capsule *in vivo*. **a)** Experimental photograph of the transanal delivery procedure for the stent-form CMU actuator and corresponding X-ray images confirming successful deployment in a porcine rectum. **b, c)** Diagram and experimental photograph of wireless charging and activation of the stent-form CMU actuator within the porcine rectum using a Tx coil. **d)** Endoscopic view of the LED switching in the stent-form CMU after activation by the outside Tx coil. **e)** Measurement of typical serum inflammatory markers before and after stent implantation in the porcine rectum. **f)** H&E staining of porcine rectal tissue samples collected from the stent site and a region distal to the stent. **g)** Diagram showing a capsule-formed MCU passing through the stent-form CMU actuator to monitor the activity of engineered bacteria within the CMU.

**Fig. S8.**
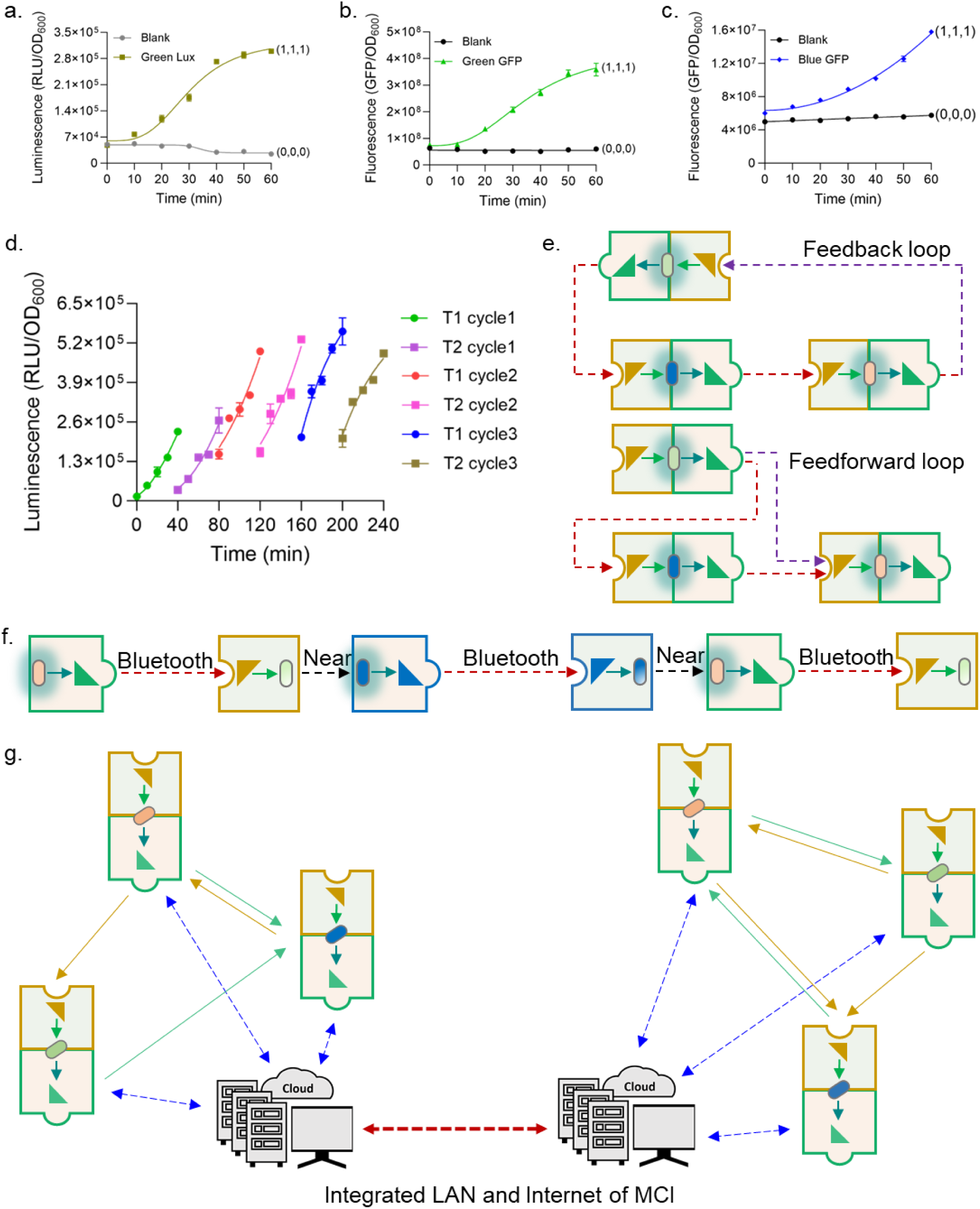
Expansion of the MCI-based communication architectures. **a, b, c)** Temporal output profiles of three receivers R1, R2 and R3 under the “1,1,1” input conditions related to the experiment in Figure 3h. **d)** Bioluminescence detection supplementary to the experiment in Figure 3l. **e)** Diagram of constructing feedback loop or feedforward loop communication with the microbe-centric transceivers. **f)** Diagram of establishing cross-life communication containing both Bluetooth-based remote communication and biochemical-based short-range communication within cellular communities. **g)** Diagram of establishing the LAN and internet of MCI.

**Fig. S9.**
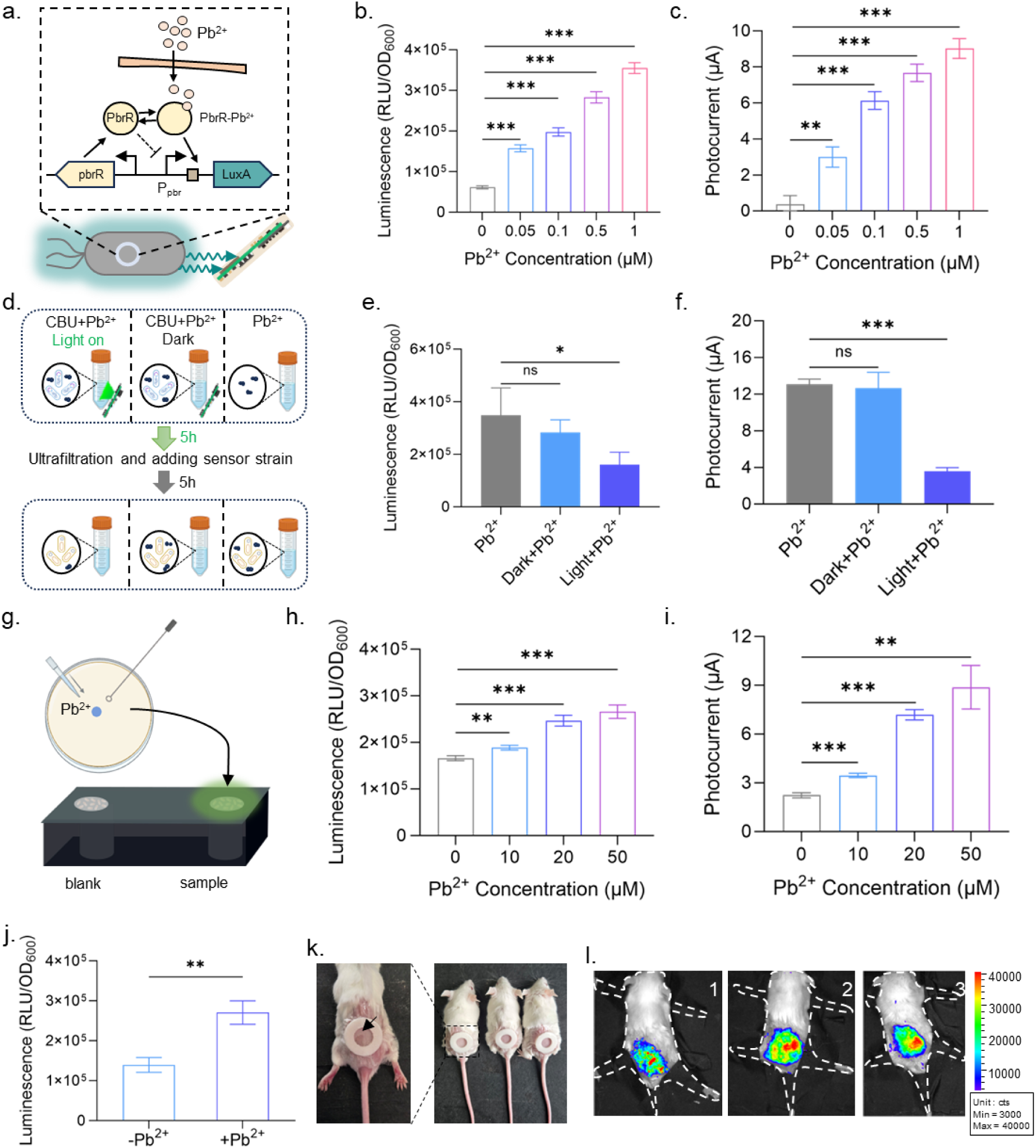
Preliminary experiment for Pb^2+^ detection *in vitro*. **a)** Genetic circuit design for Pb²⁺ sensing by the engineered bacteria within the box-form MCU sensor. **b)** Detection of various concentrations of Pb²⁺ by the sensing strain using a microplate reader. **c)** Measurement of Pb²⁺ levels at different concentrations using the MCU sensor. **d)** Experimental diagram of the opto-activated microbial expression and secretion of MtnB for Pb²⁺ chelation and removal. **e)** Evaluation of Pb²⁺ chelation efficiency described in (d) using a microplate reader. **f)** Assessment of Pb²⁺ chelation effect from (d) with the MCU sensor. **g)** Diagram showing detection of Pb²⁺ at varying concentrations on a Petri dish using the MCU sensor. **h)** Detection of Pb²⁺ in the setup from (g) using a microplate reader. **i)** Sensing of Pb²⁺ in the setup from (g) with the MCU sensor. **j)** Measurement of Pb²⁺ applied on mouse skin surface via microplate reader. **k)** Images of annular patches comprising hydrogel dressings loaded with opto-controlled EcN bacteria sealed with a polyurethane membrane for attaching to mouse skin. **l)** Fluorescence image of mice epidermis 5 hours after activating mCherry reporter secretion via the wearable CMU.

**Fig. S10.**
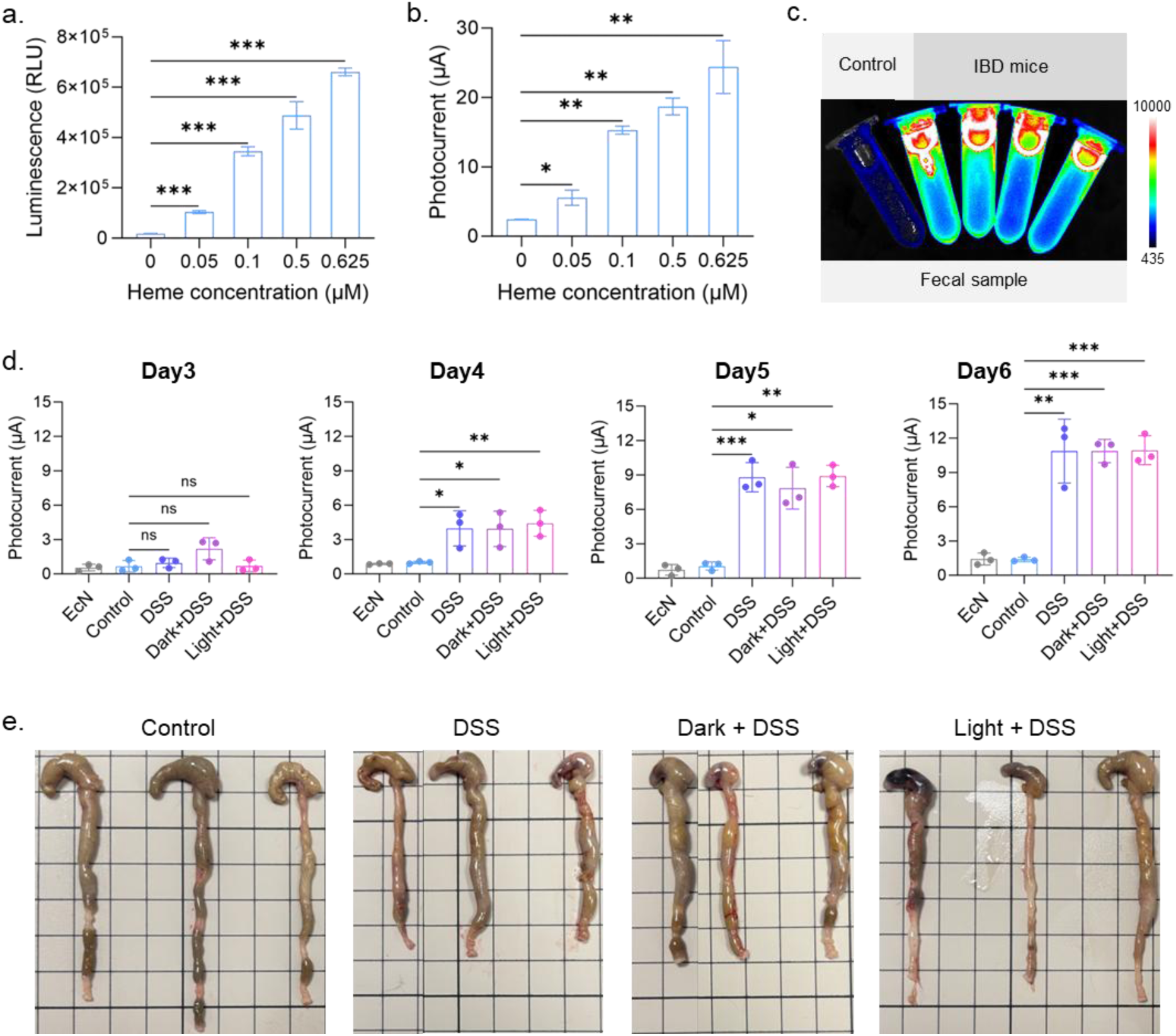
Supplemental experiments of the mouse IBD monitoring and intervention. **a)** Characterization of the response of heme-sensing bacteria to varying concentrations of heme using a microplate reader. **b)** Detection of different heme concentrations with the box-form CMU sensor. **c)** Imaging results of heme-sensing bacteria mixed with fecal samples from IBD model mice. **d)** Monitoring of fecal heme levels using the MCU sensor at different time points of the IBD mouse model timeline. **e)** Colon length measurement in mice from different experimental groups at the endpoint of the timeline of 12 days.

**Fig. S11.**
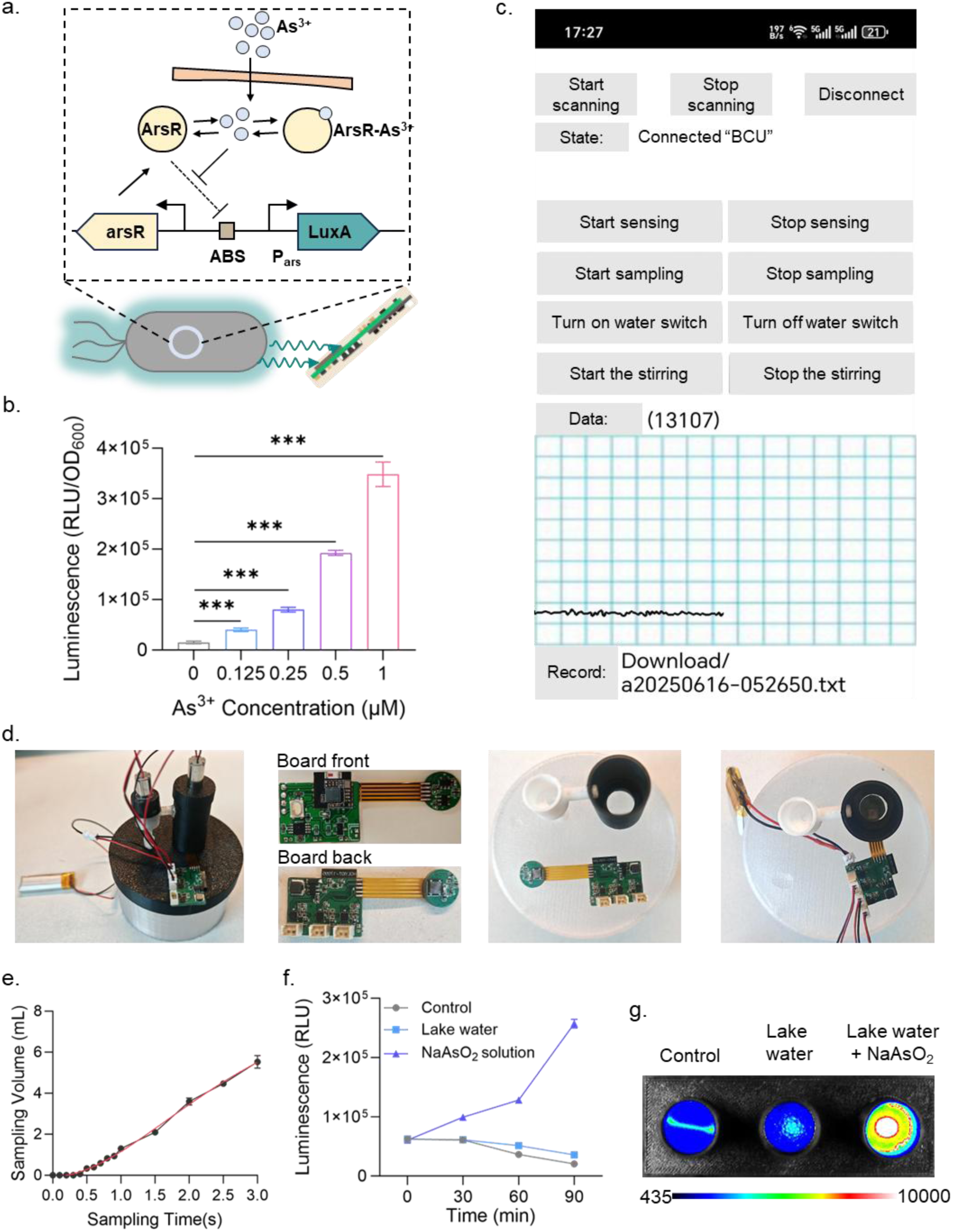
Supplemental experiments of aqueous monitoring with CMU sensor. **a)** Genetic circuit design for As³⁺/AsO₂⁻ sensing by the engineered bacteria within the box-form MCU sensor. **b)** Response of the sensor bacteria to varying concentrations of AsO₂⁻ measured using a microplate reader. **c)** App interface for controlling the MCU sensor integrated into a USV (unmanned surface vehicle) module. **d)** Images of the whole MCU sensor and therein components, including the electronic circuit board, light-shielded sampling chamber containing As³⁺/AsO₂⁻-sensing bacteria, and auxiliary wiring. **e)** Variation of the sampling volume along with time. **f)** Spiking experiment: detection of AsO₂⁻ in lake water samples using a microplate reader. **g)** Imaging results of the engineered bacterial sensor responding to AsO₂⁻-spiked lake water samples.

**Table S1.** The engineered microbial strains in this study.

| Name | Key characteristics |
| --- | --- |
| EcN | *E. coli* Nissle 1917 ΔpMUT1 ΔpMUT2 (cryptic plasmids) |
| *B. subtilis* 168 | ATCC 23857, Ind<sup>-</sup> Tyr<sup>+</sup> |
| *S. cerevisiae* BY4741 | MATa his3Δ1 leu2Δ0 met15Δ0 ura3Δ0 |
| EcN-CSG-Lux | EcN Δ*csgA*, Δ*csgD*::pJ23101-*csgEFG*, pJ23100-*csgBC*, Δ*lacZ*::pJ23100-*LuxBCDE* |
| DH5α-Lux | DH5α harboring pCons-LuxABCDE |
| *B. subtilis*-Lux | *B. subtilis* 168 integrated with Ppen-*luxABCDE* cistron at *wprA* site |
| *S. cerevisiae* BY4741-Lux | *S. cerevisiae* BY4741 transformed with pRS425 plasmid inserted by P<sub>TEF1</sub>-nanoluc-T<sub>GPM1</sub> |
| DH5α-Ara | DH5α transformed with pAra-LuxABCDE |
| EcN-Nitrate | EcN-CSG-Lux transformed with pNitrate-LuxA |
| EcN-Tet | EcN-CSG-Lux transformed with pTet-LuxA |
| EcN-Heme | EcN-CSG-Lux transformed with pHeme-LuxA |
| DH5α-Hypoxia | DH5α transformed with pHypoxia-LuxCDABE |
| EcN-Pb | EcN-CSG-Lux transformed with pPb-LuxA |
| EcN-As | EcN-CSG-Lux transformed with pAs-LuxA |
| EcN-Opto-GFP | EcN harboring p15A-*ccaS*-PCB and pGreen-sfGFP |
| EcN-Opto-Lux | EcN-CSG-Lux harboring p15A-*ccaS*-PCB and pGreen-LuxA |
| EcN-Opto-secGFP | EcN harboring pHlyBD, p15A-*ccaS*-PCB and pGreen-sfGFP-H60 |
| EcN-Opto-secCherry | EcN harboring pHlyBD, p15A-*ccaS*-PCB and pGreen-mCherry-H60 |
| EcN-Opto-secPD-L1nb | EcN harboring pHlyBD, p15A-*ccaS*-PCB and pGreen-Nb-PD-L1nb-H60 |
| EcN-Opto-secMtnB | EcN harboring pHlyBD, p15A-*ccaS*-PCB and pGreen-Nb-MtnB-H60 |
| EcN-Opto-secIL-10 | EcN harboring pHlyBD, p15A-*ccaS*-PCB and pGreen-Nb-IL-10-H60 |
| BY4741-Opto-secAmylase | *S. cerevisiae* BY4741 harboring pBlue-Amylase |

**Table S2.**
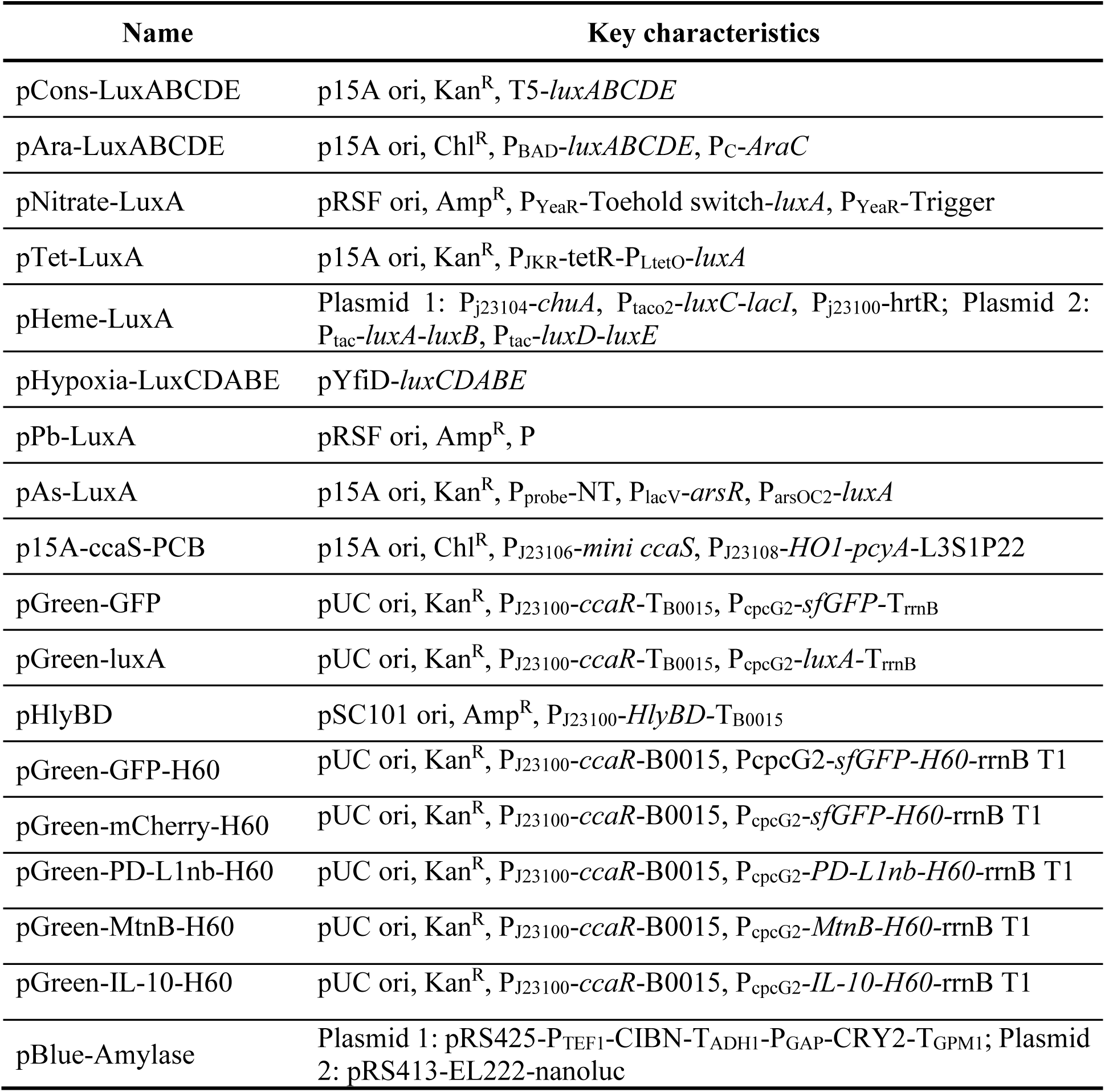
The constructed plasmids in this study.

